# Positional information drives distinct traits in transcriptomically identified neuronal types

**DOI:** 10.1101/2024.09.15.613138

**Authors:** Inbal Shainer, Johannes M. Kappel, Eva Laurell, Joseph C. Donovan, Martin W. Schneider, Enrico Kuehn, Irene Arnold-Ammer, Manuel Stemmer, Johannes Larsch, Herwig Baier

**Affiliations:** Max Planck Institute for Biological Intelligence, Am Klopferspitz 18, 82152 Martinsried, Germany; Friedrich Miescher Institute for Biomedical Research, Maulbeerstrasse 66, CH-4058 Basel, Switzerland; Center for Integrative Genomics, Faculty of Biology and Medicine, University of Lausanne, CH-1015 Lausanne, Switzerland

**Author notes:** These authors contributed equally to this work (Inbal Shainer, Johannes M. Kappel).

## Abstract

Neuronal phenotypic traits such as morphology, connectivity, and function are dictated, to a large extent, by a specific combination of differentially expressed genes. Clusters of neurons in transcriptomic space correspond to distinct cell types and in some cases (e. g., *C. elegans* neurons^1^ and retinal ganglion cells^2–4^) have been shown to share morphology and function. The zebrafish optic tectum is composed of a spatial array of neurons that transforms visual inputs into motor outputs. While the visuotopic map is continuous, subregions of the tectum are functionally specialized^5,6^. To uncover the cell-type architecture of the tectum, we transcriptionally profiled its neurons, revealing more than 60 cell types that are organized in distinct anatomical layers. We then measured the visual responses of thousands of tectal neurons by two-photon calcium imaging and matched them with their transcriptional profile. Furthermore, we characterized the morphologies of transcriptionally identified neurons using specific transgenic lines. Surprisingly, we found that neurons that are transcriptionally similar can diverge functionally and morphologically. Incorporating the spatial coordinates of neurons within the tectal volume revealed functionally and morphologically defined anatomical subclusters within individual transcriptomic clusters. Our findings demonstrate that extrinsic, position-dependent factors expand the phenotypic repertoire of genetically similar neurons.

## Main

Neurons can be grouped into classes, types, and subtypes by the combination of genes they express^7–9^. The cell type-specific transcriptome (t-type) is believed to encode the genetic instructions for a neuron’s differentiation trajectory during development and thus its morphology (m-type), connectivity and function (f-type). For example, each of the 118 anatomically distinct neuron types in the roundworm *C. elegans* expresses a unique, sparse combination of transcription factors, which regulate downstream genes, thus shaping the neuron’s phenotype and contribution to network function^1,10^. In *Drosophila*, distinct sets of differentially expressed genes are associated with specific axonal- or dendritic-wiring^11^. Similarly, in the mouse visual cortex, morpho-electric properties measured in tissue slices were found to be relatively homogeneous within individual t-types^12,13^. However, the dogmatic view of “t-type = m-type = f-type” is problematic, as functional responses and dendritic arbor elaborations are often shaped by modulatory influences and individual experience^14,15^. For instance, GABAergic interneurons in the visual cortex were found to exhibit tuning properties dependent on behavioral state rather than t-type^16^. Moreover, mouse cortical neurons of the same t-type may vary strongly in their functional tuning^17^, as well as in their m-type, showing divergent long range projections and local connectivity^18^. It is therefore essential to determine the extrinsic factors that influence the expression of a neuron’s phenotype and how these interact with the transcriptome.

In this study, we provide evidence that the interplay of transcriptome, development, and topography shapes neuronal phenotypes. The zebrafish optic tectum (OT) receives topographically organized input from retinal ganglion cells (RGCs) along the anterior-posterior (AP) and dorsal-ventral (DV) axes^19,20^. In its superficial-to-deep (SD) dimension, orthogonal to the retinotopic axes, the OT contains a neuropil layer, in which RGC axons form synapses with the dendrites of tectal neurons, and a cell-body layer, the stratum periventriculare (SPV). Cellular birthdating studies have shown that newborn neurons are initially added radially to the OT from the periventricular zone and later to the posterior margins^21,22^, with older neurons being gradually displaced into the SPV as they mature and extend neurites into the neuropil^23,24^. Extrinsic factors, including morphogens, chemotropic and cell-surface factors, vary over the course of development and along all three axes of the OT^25^. These cues might affect the neurons’ phenotypic traits in a location-dependent manner. We show here that a combination of both transcriptomic identity and cell-body position are determinants of a neuron’s phenotype in the OT.

### Diversity of neuronal and non-neuronal tectal cell types

To characterize the t-type composition of the zebrafish OT, we performed droplet-based single-cell RNA sequencing (scRNA-seq; see Methods; Extended data Fig. 1) of the OT of 6-7 dpf larvae. We sequenced 45,766 cells, which corresponds to >7x coverage of cells in a single tectal hemisphere (∼5,800 cells ^5^), and grouped them according to their transcriptomes into 25 major clusters (Fig. 1a, Extended data Fig.1). Based on differentially expressed (DE), cluster-specific genes (Supplementary Table 1, Extended data Fig. 2-3), we identified three clusters of progenitor cells, one cluster of radial glia, fourteen clusters of neurons and six additional non-neuronal populations (Extended data Fig. 2-3).

**Fig. 1.**
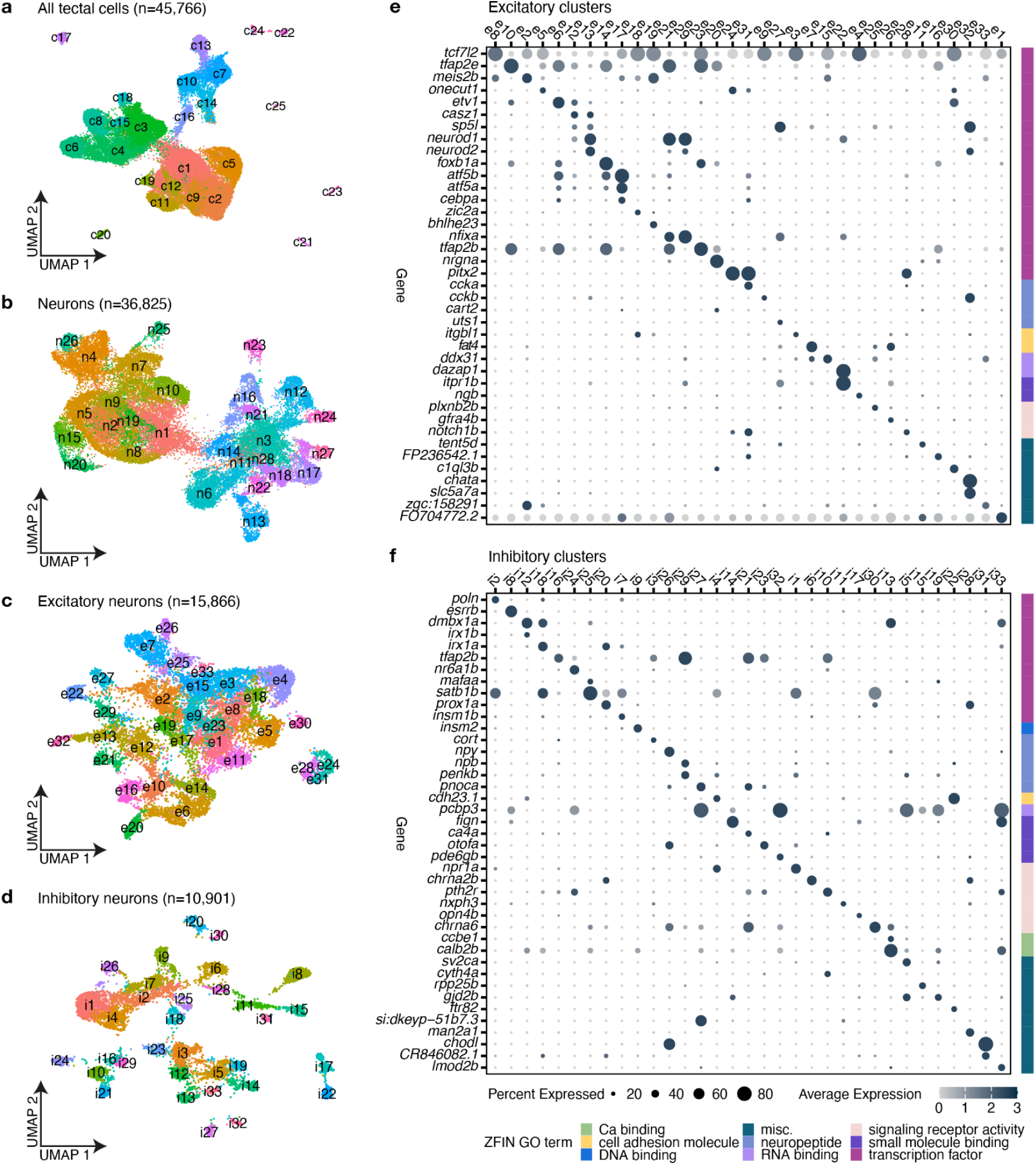
scRNA-Seq of the optic tectum reveals a multitude of neuronal types. **a.** All sequenced tectal cells clustered according to similarity of gene expression with 25 distinct clusters identified. Each dot represents a single cell, color coded according to cluster identity. **b.** Postmitotic neurons (clusters expressing *elavl3*) were subsetted and reclustered to further identify the various types of excitatory and inhibitory neurons.. **c.** Excitatory neurons reclustered according to similarity of gene expression. **d.** Inhibitory neurons reclustered according to similarity of gene expression. **e.** Dotplot of the highly DE genes for each of the excitatory cell types (clusters: e1-e33). DE genes were grouped according to their molecular function and annotated according to ZFIN GO terminology^27^. **f.** Dotplot of the highly DE genes for each of the inhibitory cell types (clusters: i1-i33).

**Extended data Fig. 1.**
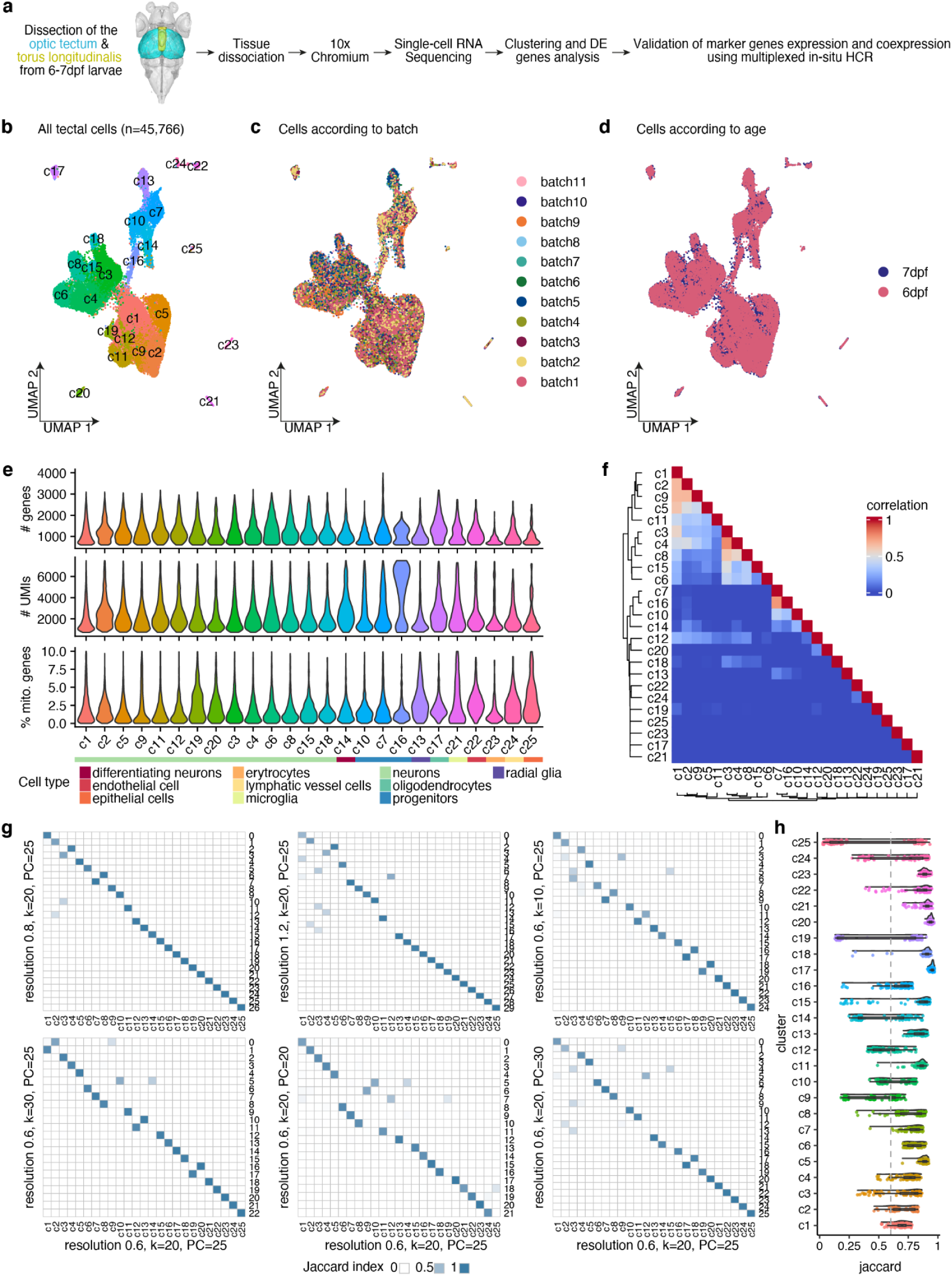
QC of the scRNA-Seq clustering analysis. **a.** Experimental procedure: optic tectum and torus longitudinalis (TL) were dissected from 6-7 dpf WT larvae in eleven experimental batches. The cells were dissociated and single-cell barcoded cDNA was generated using the 10x Chromium system and sequenced. Cells were clustered into t-types according to the similarity of gene expression. **b.** After batch correction using Harmony^28^ and removal of immediate early genes (see Methods), we identified a total of 25 clusters. Each dot represents a single cell, color coded according to the cluster. **c.** Same UMAP as in **b**, color coded according to the batch. Clusters include cells from all batches. **d.** Same UMAP as in **b**, color coded according to the larvae age. **e.** QC metric of the sequenced cells after filtration, according to cluster. Cells were filtered (see Methods) in order to remove outliers that contain either low or high number of genes and UMI (representing either poorly sequenced cells or doublets), as well as cells containing high percentage of mitochondrial genes (representing stressed cells). Different cell types contained different levels of genes and UMIs, matching their biological profile (for example high levels of UMIs in some of the progenitor cells). **f.** The similarity between clusters was assessed by correlating the specificity scores of the top markers associated with each cluster (see Methods). The vast majority of clusters showed low correlation with other clusters, representing unique t-types. **g.** The robustness of the clusters was assessed by measuring the overlap of cells grouped together under different clustering parameters in comparison to the parameters used (see Methods). **h.** Jaccard raincloud plot showing cluster stability. 80% of the data was repeatedly subsampled, and the pairwise Jaccard index was measured for each cluster before and after subsampling and reclustring (see Methods). In most instances, we observed that clusters with small numbers of cells were scored low, as a result of infrequent sampling of those cells.

**Extended data Fig. 2.**
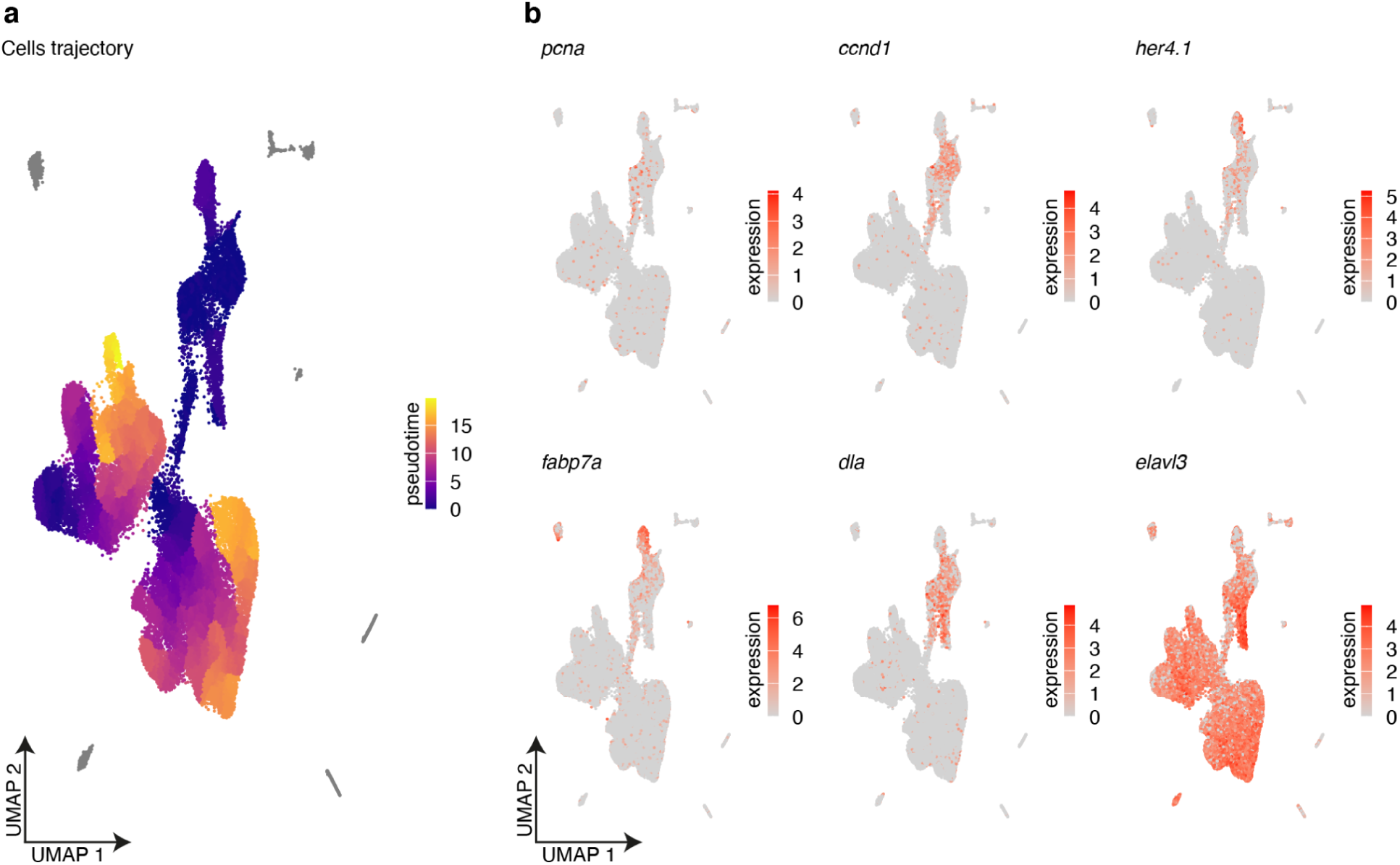
Pseudotime analysis. **a.** Pseudotime analysis was performed with Monocle3^29^. Cells expressing proliferating cell nuclear antigen (*pcna*), a key component of the DNA replication machinery and a marker of the G1 phase^30^, and belonging to clusters c7, c10, and c16, were identified as progenitor cells and defined as the root point. Cells are color coded according to their pseudotemporal ordering. **b.** Expression patterns of selected marker genes, representing the different stages of proliferation and differentiation. The marker *ccnd1* is a member of the cyclin family that regulates the cell-cycle transition from G1 to S phase ^31^. The markers *fabp7a* and *her4.1* are typically expressed in radial glia^32,33^. Delta A (*dla*) regulates cell fate specification^34^, and *elavl3* is expressed in committed neurons.

**Extended data Fig. 3.**
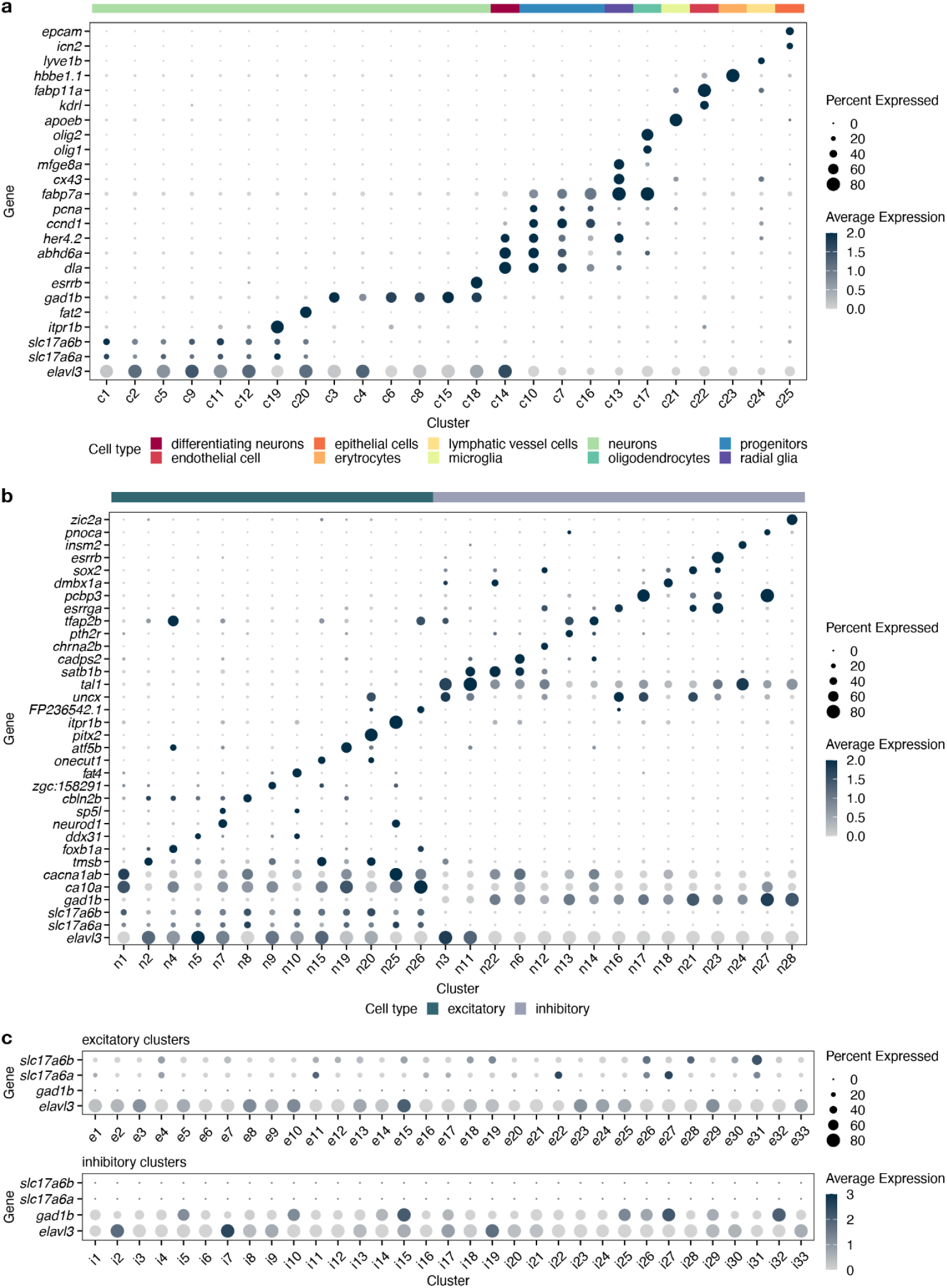
Cell type classification. **a.** Dotplot of the highly DE genes for each tectal cluster. Clusters were grouped according to cell function. Cluster c20 cells expressed torus longitudinalis marker genes and were therefore omitted from downstream tectum analyses. **b.** Dotplot of the highly DE genes for each neuronal cluster. Clusters were grouped according to transmitter use and developmental stage. Cluster n3 highly expresses the transcription factors *uncx* and *tal1,* which are typically expressed in differentiating neurons^35–37^. **c.** Based on *gad1b*, *slc17a6a* (*vglut2b*) and *slc17a6b* (*vglut2a*) expression, we separated the inhibitory and excitatory clusters and reclustered them (see Methods).

**Extended data Fig. 4.**
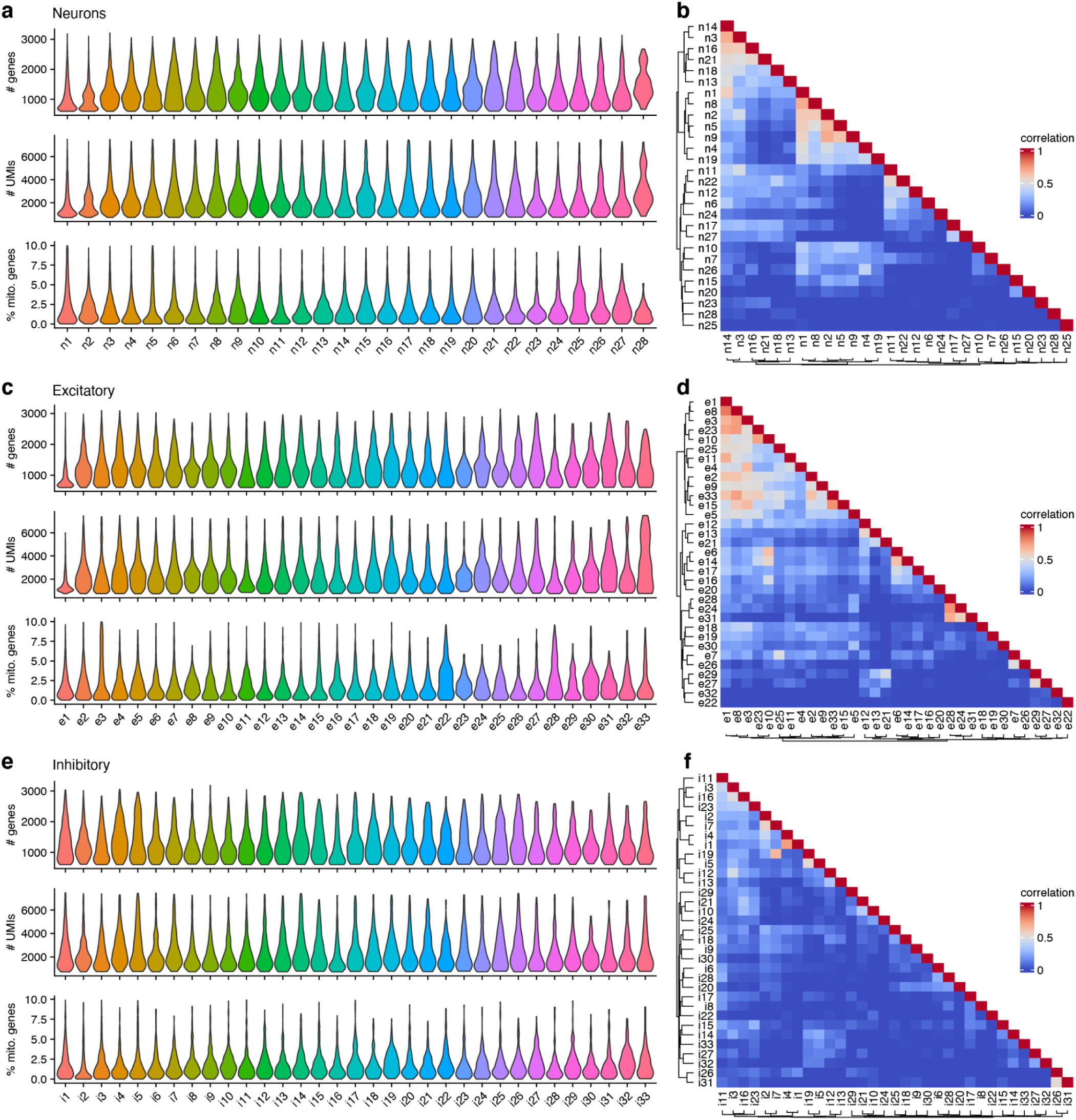
QC metrics for clustering of excitatory and inhibitory cells. **a.** Distribution of gene count, UMI count and percentage of mitochondrial genes per neuronal cluster. **b.** The similarity between neuronal clusters was assessed by correlating the specificity scores of the top markers associated with that cluster (see Methods). **c.** Same as **a,** per excitatory cluster. **d.** Same as **b**, for the excitatory clusters. **e.** Same as **a** per inhibitory cluster. **f.** Same as **b**, for the inhibitory clusters.

**Extended data Fig. 5.**
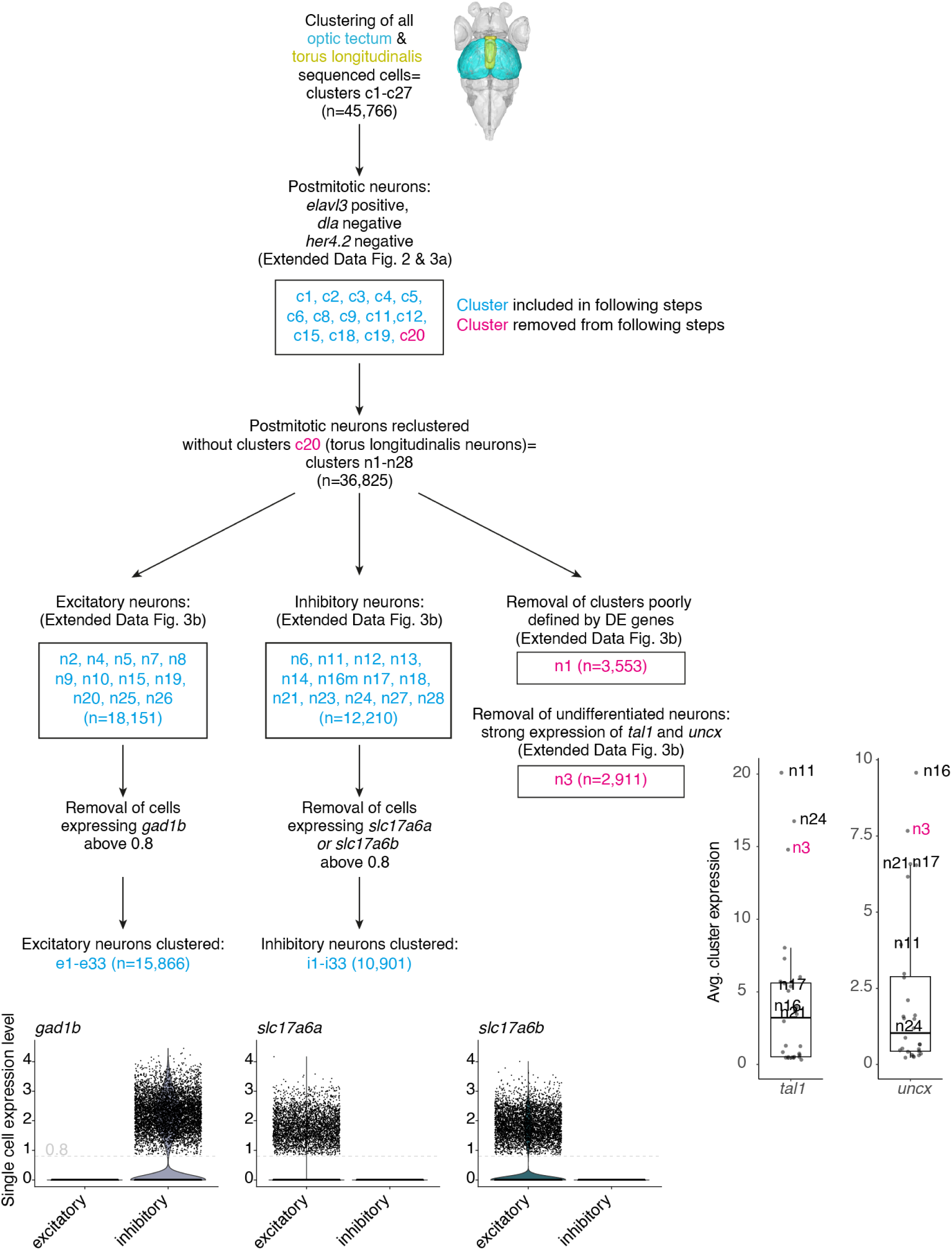
Summary of scRNA-seq clustering steps.

**Fig. 2.**
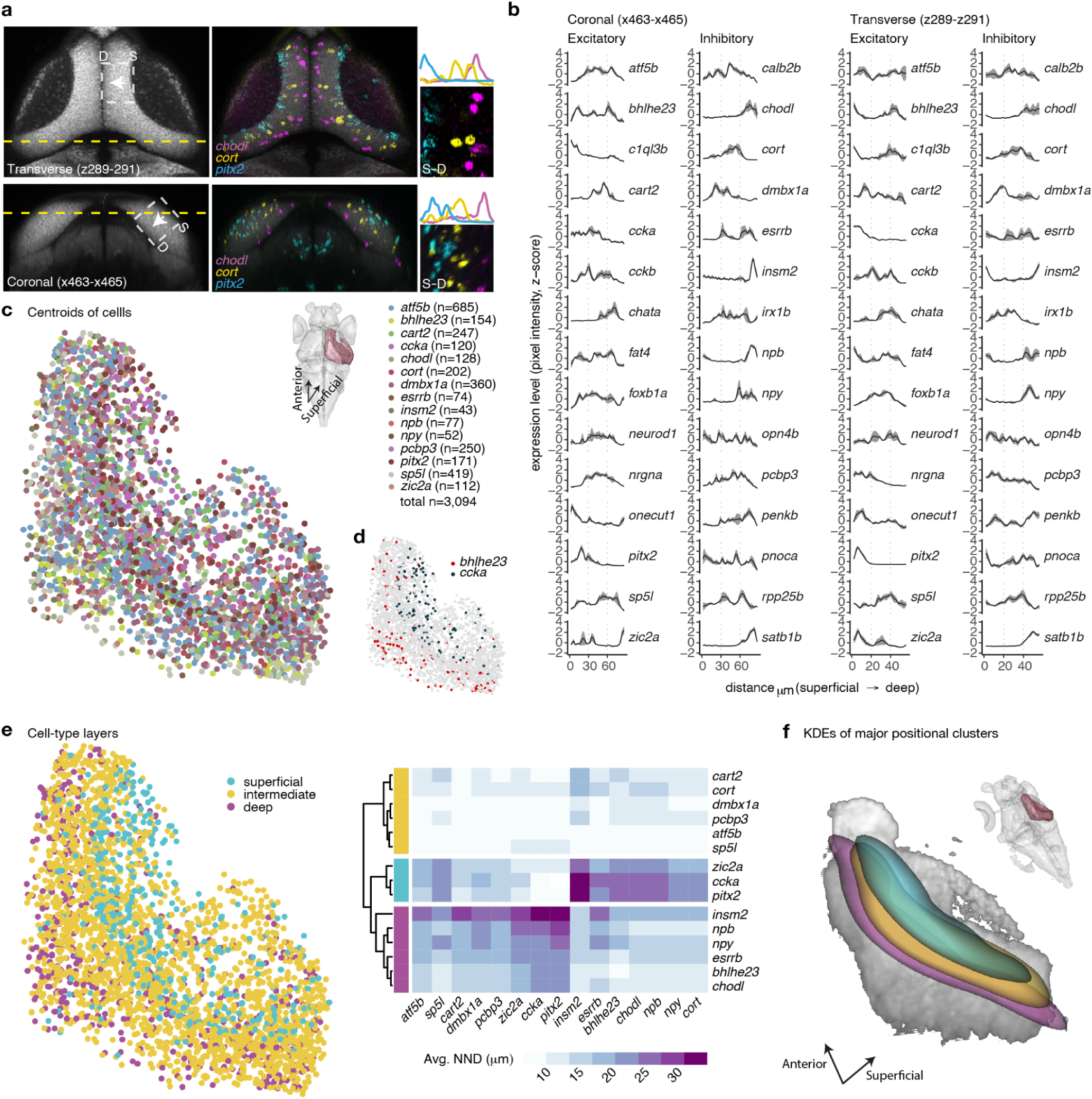
Transcriptomic cell types form molecular layers orthogonally to the retinotopic map. **a.** Multiplexed in situ HCR of selected marker genes were registered to the reference brain of the mapzebrain.org atlas. We measured the expression level by examining the pixel intensity profile of each gene in the same transverse (projection of planes z289-z291) and coronal section (projection of planes x463-x465). The pixel intensity measured area is labeled with a white dashed rectangle and confined to the SPV (S; superficial, D; deep). **b.** Z-scored pixel intensity of selected inhibitory and excitatory marker genes. Black line represents the average (n=3, simple moving average with window size=3), the gray shade represents standard error. **c.** Centroids of selected tectal cell type markers located within the SPV were manually labeled. Dorsal view with the right hemisphere tectal SPV highlighted in burgundy is shown, representing the labeled area of the centroids, as well as the tectal coordinate system. **d.** The labeled centroids of *bhlhe23* and *ccka* are shown, demonstrating the segregation of these cells along the tectum in 3D. **e.** The distance from each cell in a given cell type to its nearest neighbors of all other cell types was measured in 3D (NND). Hierarchical clustering of the average NND for each cell type revealed three main clusters, dividing the SPV layer into three molecular layers. **f.** 3D visualization of thresholded Gaussian kernel densities for the three molecular layers from **e.** Inset shows the position of this region within the whole brain.

**Extended data Fig. 6.**
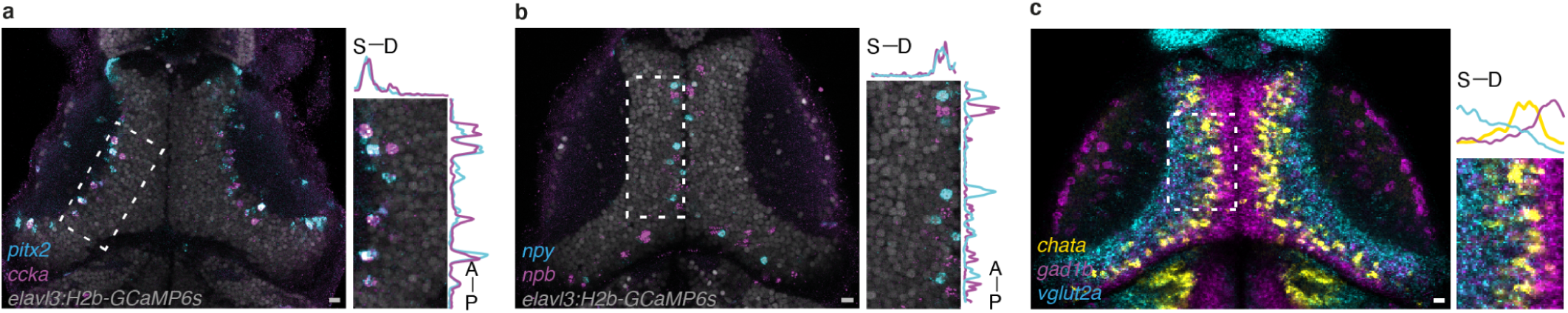
Multiplexed in situ HCR labeling of cell type markers. We applied in situ HCR to analyze and validate the spatial arrangement of the cell types’ marker genes. **a.** The marker gene *ccka* was co-expressed with *pitx2* in cluster e31 but not in cluster e24. Dashed square area is enlarged and the pixel intensity (z-scored) is plotted, demonstrating both the co-expression and the localization to the superficial part of the SPV (S; superficial, D; deep). Scale bar = 10 µm. **b.** The expression of the neuropeptides *npb,* marker of clusters i22 and i29, and *npy,* marker of cluster i26 was mutually exclusive. In situ HCR analysis demonstrated the localization to the deep sublayer of the SPV. Scale bar = 10 µm. **c.** Registered in situ data of *gad1b*, *vglut2a* (*slc17a6b*), and *chata* revealed that glutamatergic, cholinergic, and GABAergic neurons are segregated along the tectal superficial to deep axis. Scale bar = 10 µm.

**Extended data Fig. 7.**
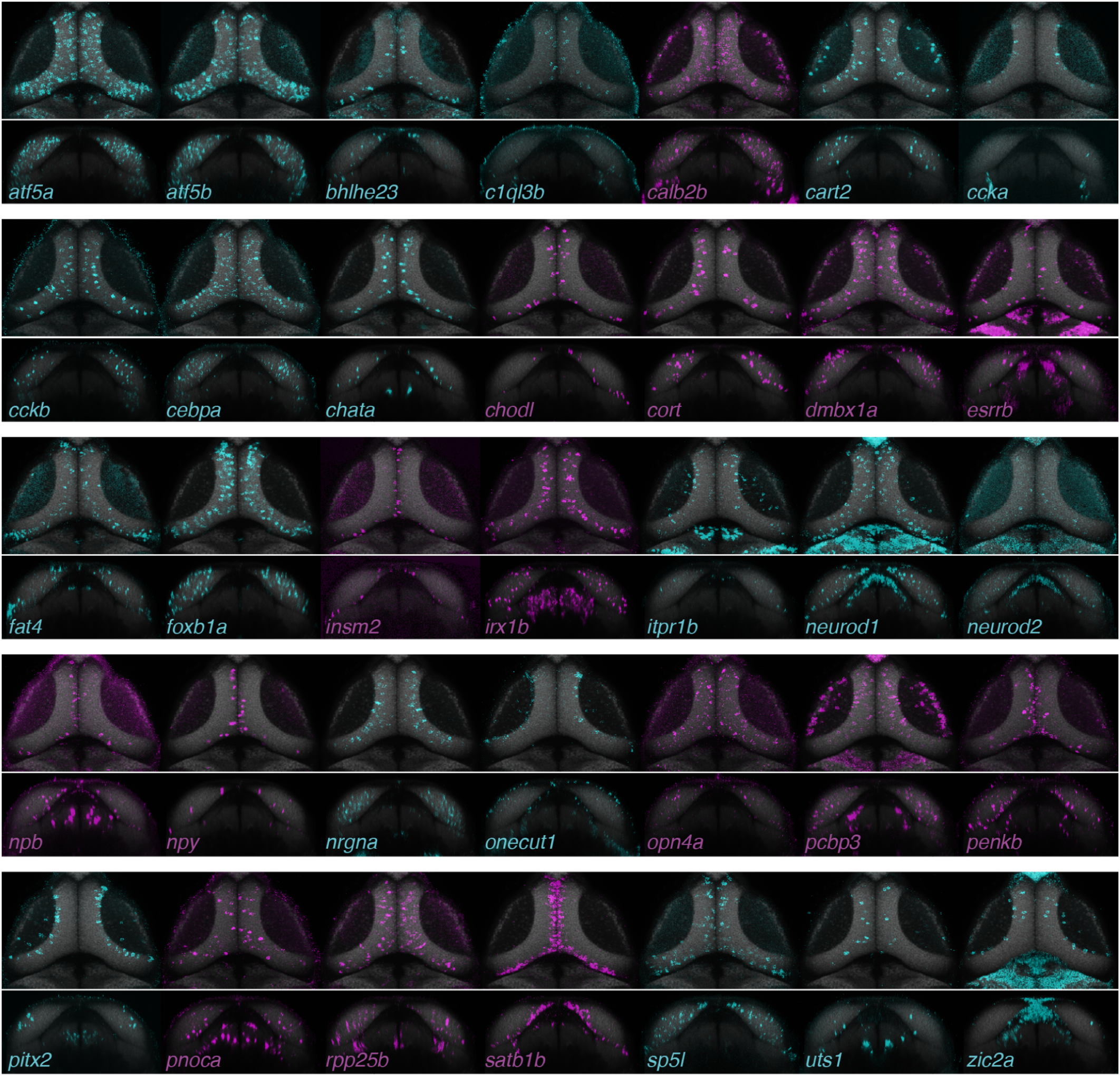
Marker genes expression patterns. Registered in situ HCR data for the examined cell type markers. Projection of transverse planes z289-z291 and coronal planes x463-x465 of single animals. Excitatory markers are shown in cyan, inhibitory markers are shown in magenta. All the data can be viewed through mapzebrain.org^39^.

**Extended data Fig. 8.**
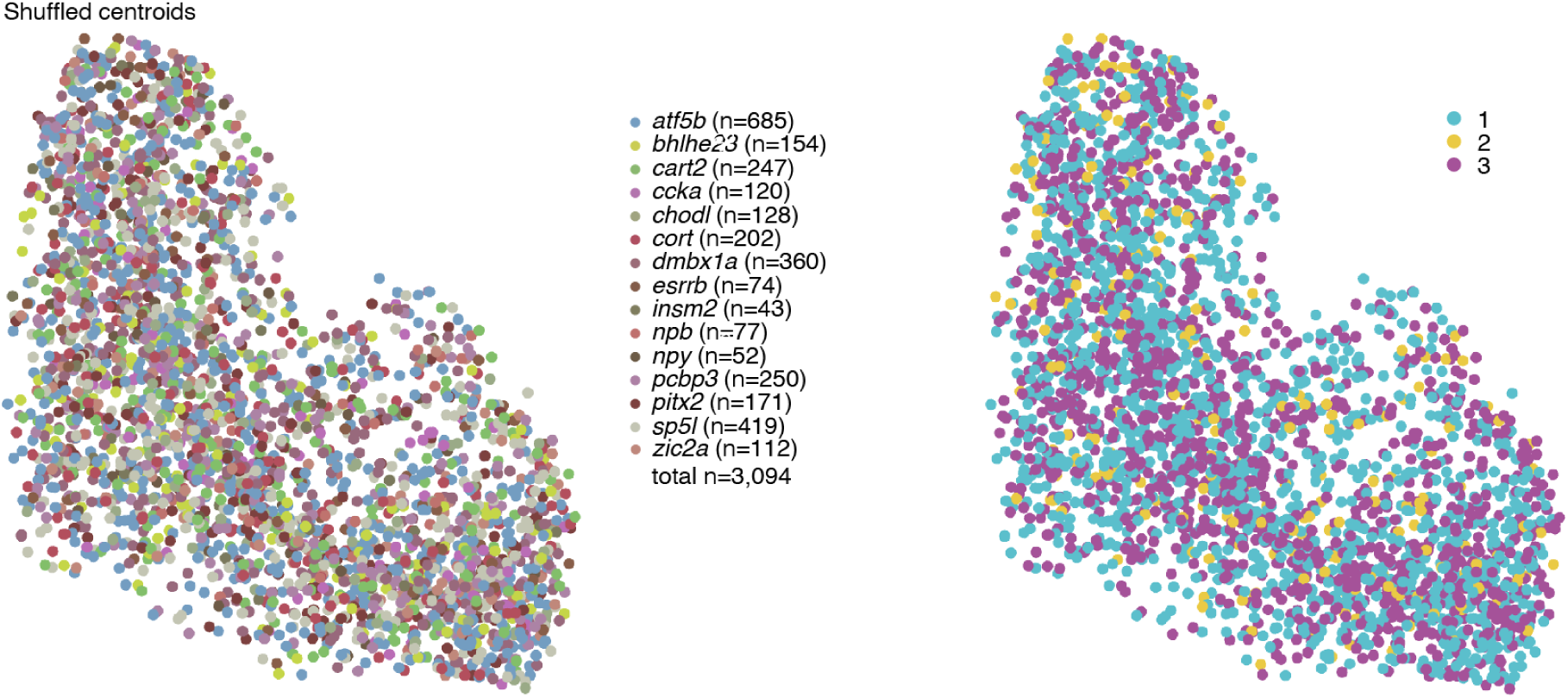
Shuffling of the tectum cell type centroid labels results in a loss of the observed molecular layers within the tectum.

**Figure 3.**
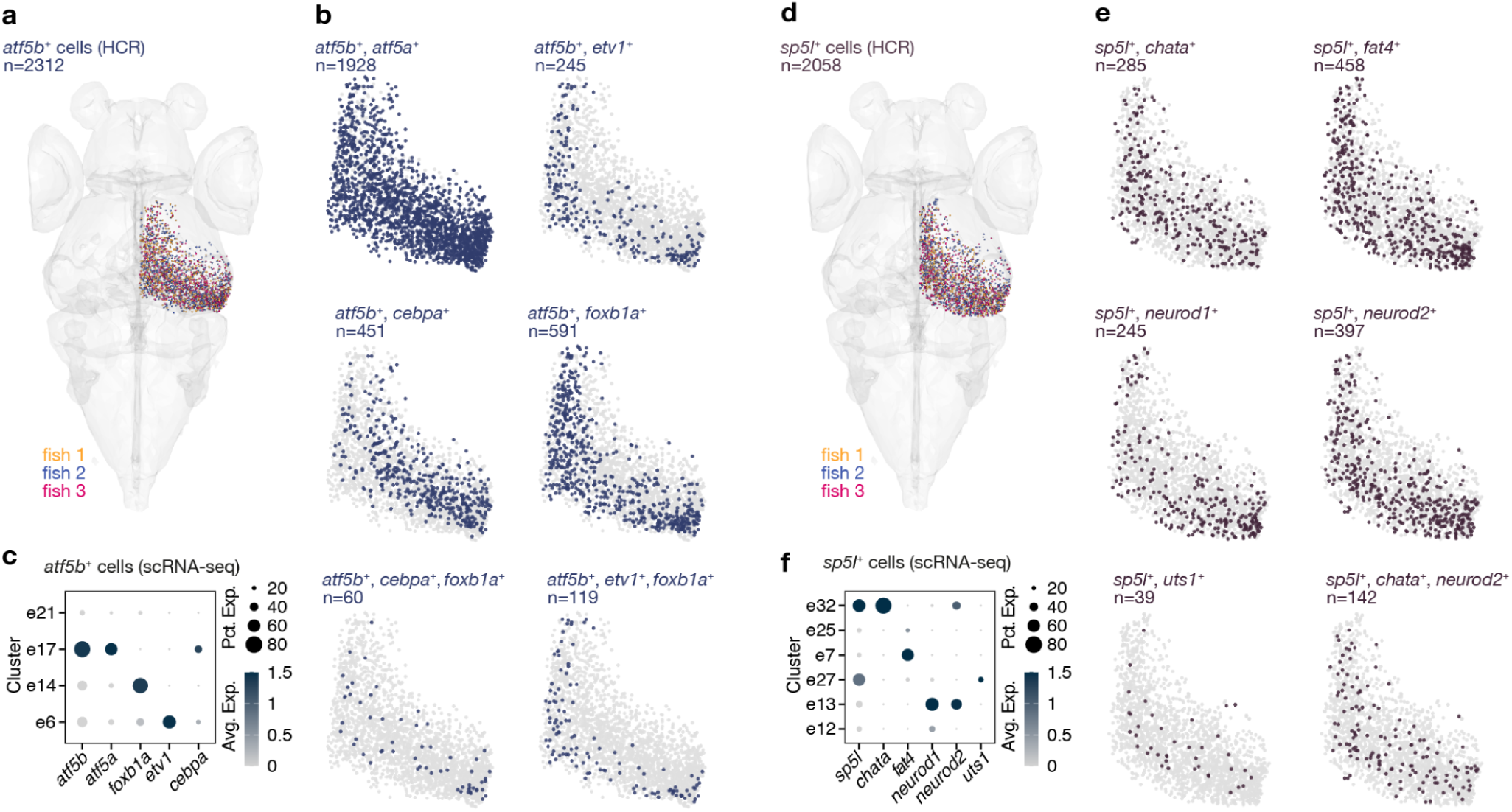
Combinatorial expression of t-type marker genes separate subtypes along the anatomical axes of the OT. **a.** *atf5b*-expressing cells were labeled from 3 individual larvae (total n=2312). **b.** Combinatorial expression of marker genes within the *atf5b*-expressing cells. *atf5b+ etv1+* cells were found in the deep SPV region, while *atf5b+ cebpa+* cells were localized to the superficial region with highest density towards the posterior zone. **c.** The scRNA-seq data of clusters expressing *atf5b*. **d.** *sp5l*-expressing cells were labeled from 3 individual larvae (total n=2058). **e.** Combinatorial expression of marker genes within the *sp5l*-expressing cells. *sp5l+ chata*+ cells were found in the intermediate SPV region. *sp5l+ neurod1+* cells were found in the intermediate and deep regions, with highest density towards the posterior zone. *sp5l+ uts1+* represented a rare population located in the intermediate SPV region. **f.** The scRNA-seq data of clusters expressing *sp5l*.

The postmitotic neurons (36,825 cells, expressing *elavl3*) were reclustered (Fig. 1b, Extended data Fig. 2-4). Poorly annotated clusters were removed (Extended data Fig. 5), and the remaining clusters were separated into excitatory (*gad1b*-, *slc17a6a*+, *slc17a6b+*) and inhibitory (*gad1b*+, *slc17a6a*-, *slc17a6b*-) neurons (Fig. 2c-d, Extended data Fig. 3c & 4). This resulted in 33 excitatory t-types (clusters: e1-e33; total number of cells = 15,866) and 33 inhibitory t-types (clusters: i1-i33; total number of cells = 10,901). For 20 neuronal types, we identified individual DE genes that serve as mutually exclusive markers; for the remaining types, sparse combinations of DE genes sufficed for an unambiguous definition (Fig. 1e-f, Supplementary Table 1). Overall, we identified 66 neuronal t-types in the larval zebrafish OT (Extended data Fig. 5). The composition of cell types and marker genes we identified differed from a recent study characterizing the zebrafish OT^26^. In that study, FACS sorting of tectal neurons from a specific enhancer trap line was used prior to the scRNA-seq, which biased the data toward the fraction of cells labeled by that line^26^. This most likely resulted in an incomplete coverage of the OT cell types and the observed higher proportion of immature neuronal populations. Our approach instead did not restrict the access to the complete range of tectal cells, which resulted in identification of a rich repertoire of neuronal and non-neuronal populations.

**Fig. 4.**
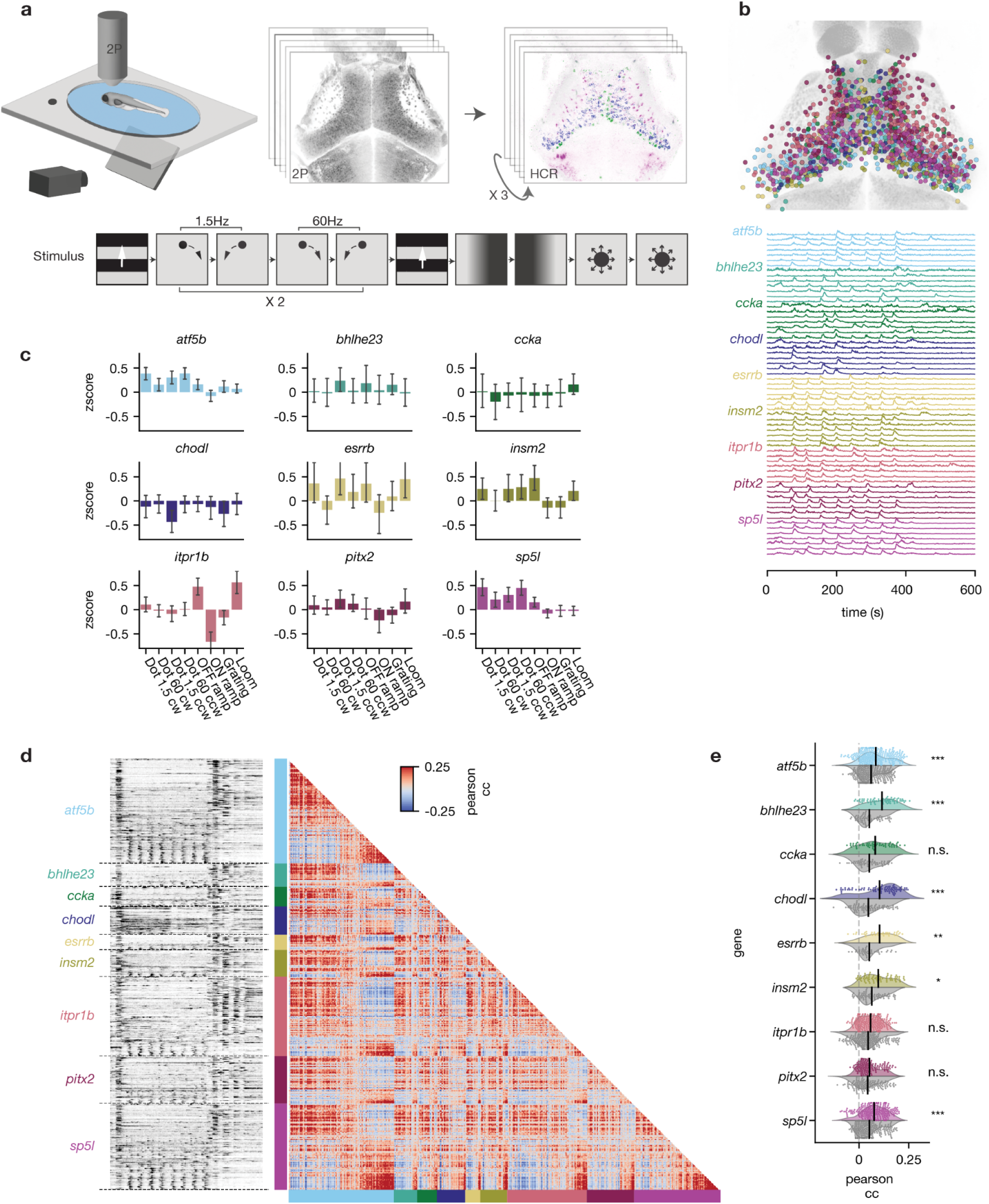
t-types show diverse visual responses and form coherent functional subclusters. **a.** Experimental procedure: 6 dpf larvae were exposed to a battery of visual stimuli while volumetric functional 2-photon calcium imaging was performed covering most neurons of the tectum. The larvae were then stained in consecutive rounds for up to 6 marker gene mRNAs using HCR labeling. **b.** Top: Result of aligning functional and HCR brain volumes and registering all ROIs that overlap with one of 9 marker genes into a common anatomical reference frame (ROIs=1204, N=6). Bottom: Example traces of each labeled t-type selected for responsiveness to local motion. **c.** Average t-type response scores to the visual stimulus sequence scaled to unit variance and zero mean of overall tectal population. Most t-types showed elevated or decreased responses to at least two different stimuli. **d.** Left: Raw calcium traces of all 1204 t-type+ functional ROIs sorted by first t-type and within t-type by response score to local motion. Responses to global motion (beginning, end) as well as local motion (middle section) are visible in all t-types. Right: Correlation matrix of all pairwise correlations using pearson correlation coefficient. Red and blue clusters of positively and negatively correlated neurons are found between and within all t-types. **e.** Mean pairwise correlations of each neuron with all neurons of the same t-type (color) and all neurons (gray) of other t-types. Within t-type correlation is significantly increased for six out of nine tested t-types. Two-sided Mann-Whitney-U Test, Bonferroni-corrected. *p < 0.05, **p < 0.01, ***p < 0.001.

**Extended data Fig. 9.**
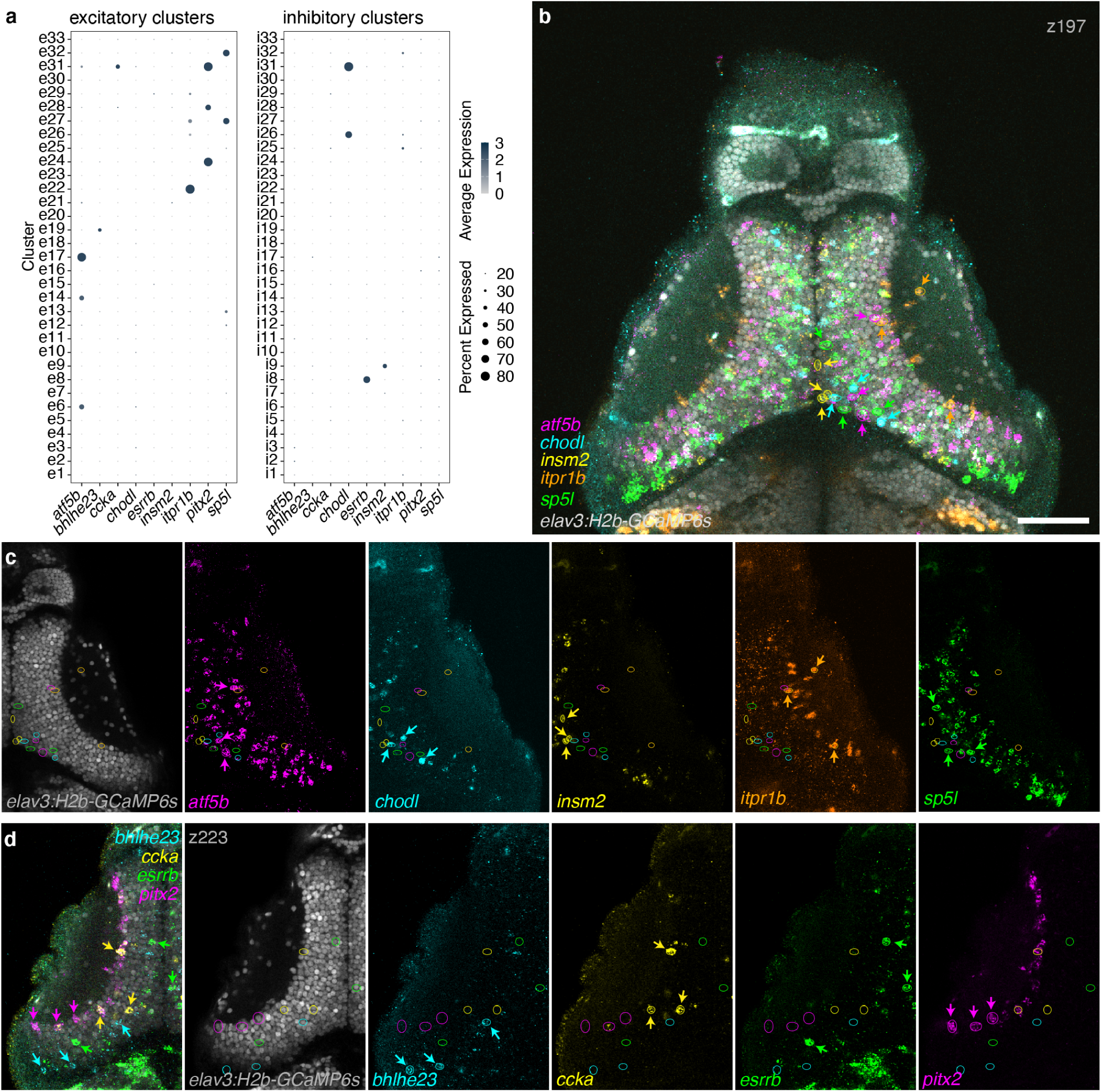
HCR labeling of marker genes post calcium imaging. **a.** HCR was performed on a set of nine marker genes. The expression levels of these genes measured by scRNA-Seq showing their expression is restricted to one or a few non-overlapping clusters, except for *ccka* that overlaps with *pitx2* expression. **b.** Multiplexed iterative HCR labeling post functional imaging. Five genes were labeled and registered. Similar to the scRNA-Seq data, these genes were not co-expressed in the same cells, representing different tectal t-type populations. Circles and arrowheads point to example cells shown in **C**, color coded similar to the gene HCR color. Single z-plane is shown, scale bar = 50 µm. **c.** Same view as **b**, splitted according to the genes labeled. **d.** Example of another multiplexed iterative HCR post functional imaging, with a different set of labeled cell type marker genes.

**Fig. 5.**
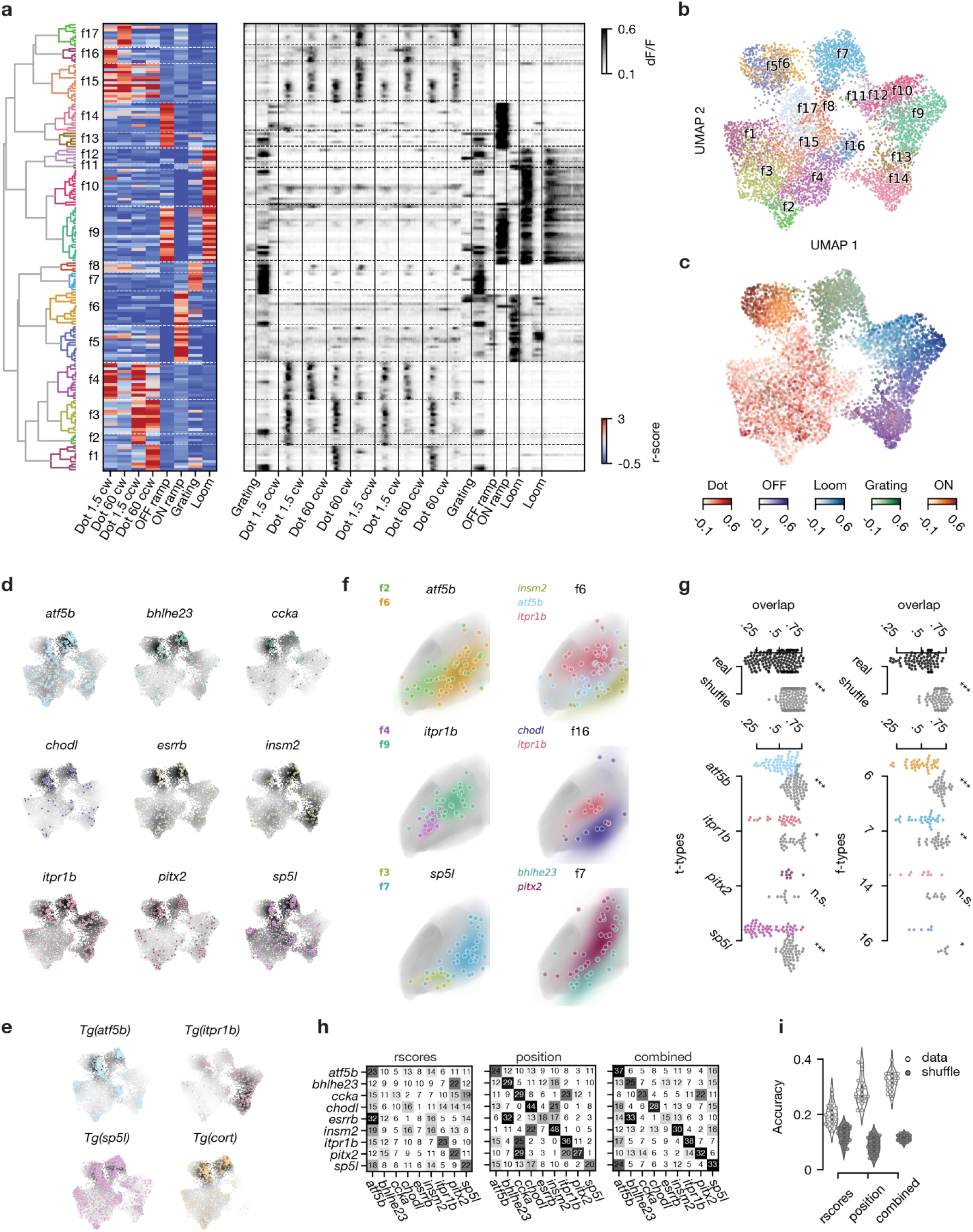
Localization in functional and anatomical space varies between t-types. **a.** Hierarchical clustering of all strongly responding tectal neurons (N=7094) to identify functional cell types (f-types). Left: Dendrogram of clustering all 169 exemplars that were identified using affinity propagation. Middle: Heatmap showing response vectors of all exemplars based on visual stimuli. Functional clusters and exemplars are primarily divided into local (clusters 1-4 & 15-17) vs. global (clusters 5-14) motion responses. Right: Raw calcium traces of all exemplars. **b.** UMAP embedding based on response vectors, overlaid with color code according to the f-type assignment. **c.** UMAP embedding of all responding neurons from **b**, with five different color codes based on the preferred respective responses of the five superclusters. **d.** t-types accumulate locally in functional space. Response vectors of each transcriptomically identified neuron transformed into the UMAP embedding. The color code of UMAP represents the kernel density estimate of the respective t-type. **e.** Response vectors of neurons recorded in transgenic animals, mapped into the same UMAP embedding largely confirm the enrichments observed in **d**. **f.** Anatomical localization of t/f-clusters within t-types and f-types. Dots represent individual ROIs, colored areas show gaussian kernel density estimates (KDEs) of t/f-clusters. ROIs of the same t-type (left) / f-type (right) comprise anatomically separated clusters based on functional (left) / transcriptomic (right) identity. All ROIs mirrored to the left tectal hemisphere. T-type is based on HCR labeling. **g.** t/f-clusters are significantly separated in anatomical tectal space within t-types and f-types. Left: Pairwise KDE overlap values for cell types of different functional clusters for real data and shuffled t-type labels. Right: Same as left for different t-types within functional clusters. Mean pairwise overlap values of t/f-clusters across t-types and across f-types are significantly lower than respective shuffled controls, respectively, indicating anatomical separation of cell types within functional clusters. Two-sided Mann-Whitney-U Test, Bonferroni-corrected. *p < 0.05, **p < 0.01, ***p < 0.001. **h.** Confusion matrices of three SVM classifiers predicting transcriptomic identity based on functional response vectors, cell body position, or both. Numbers and saturation indicate the true-positive rate. Predictive performance increases from left to right, indicated by the saturation of the diagonal (true positive predictions per cell type). **i.** Accuracy of SVM classifier performances from **h** (grey dots and violin plots). Classifiers could recover t-type identity based on position in functional space in about one out of five cases, for anatomical space accuracy was 10% higher. Combining both spaces resulted in elevated performance. For all classifiers, negative controls with shuffled cell type labels resulted in significantly lower performance (white dots and violin plots).

**Extended Data Fig. 10.**
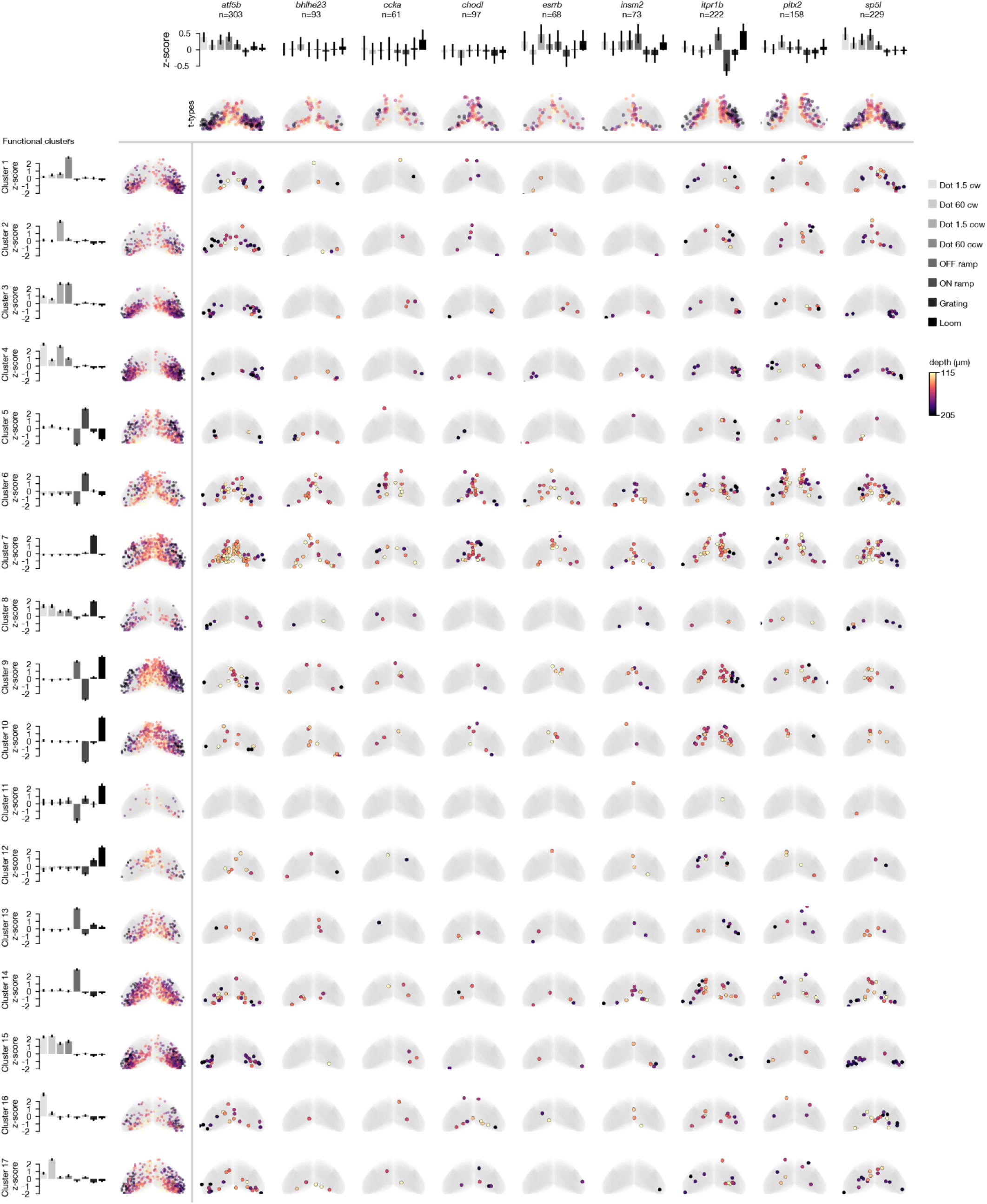
t-types comprise anatomically localized functional clusters. Left: average response scores of strongly responding neurons of each f-type scaled to tectal population mean (color coded according to stimulus type) along with their anatomical distribution. Top: average response scores of HCR labeled neurons for each t-type scaled to tectal population mean (color coded according to stimulus type) along with their anatomical distribution. Inner matrix: Anatomical localization of HCR labeled and recorded neurons separated into assigned f-types (rows) and t-types (columns). Color code depicts tissue depth.

**Extended data Fig. 11.**
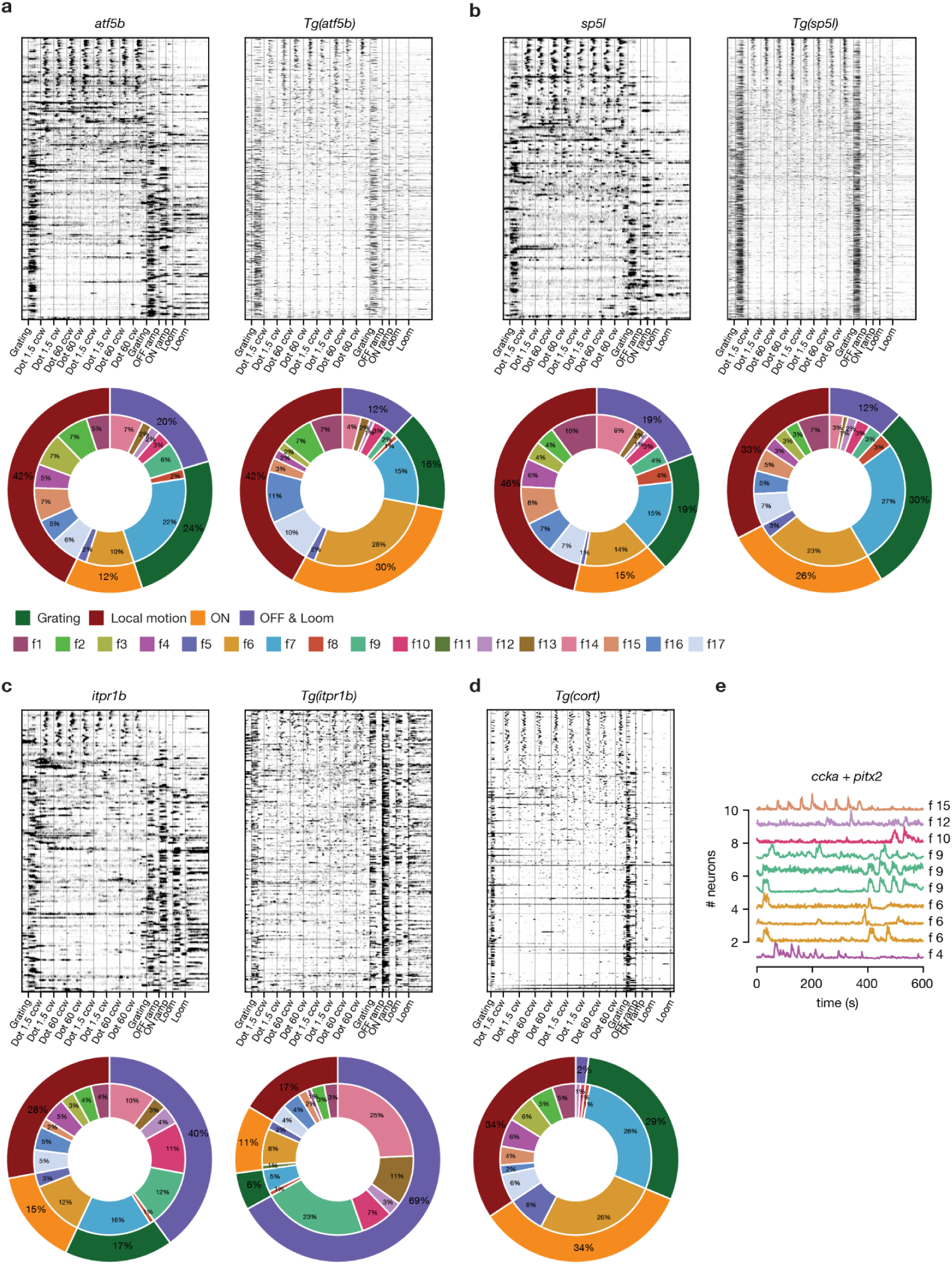
Comparison of functional responses between HCR labeled t-types and the corresponding transgenic lines. **a.** Top: Raw calcium traces of HCR labeled (left, n=303) and transgenic (right, n=1211) *atf5b+* neurons, sorted by overall response score to local motion in descending order. Bottom: Relative fractions of functional types (inner ring, according to Fig. 5a) and groups (outer ring, according to Fig. 5c) within *atf5b*+ neurons for HCR labeled (left) and transgenic (right) cells. **b.** Same as **a**, but for HCR labeled (left, n=229) and transgenic (right, n=2250) *sp5l*+ neurons. **c.** Same as **a**, but for HCR labeled (left, n=222) and transgenic (right, n=332) *itpr1b*+ neurons. **d.** Same as **a**, but for transgenic *cort*+ neurons (n=450). **e.** Raw traces and functional types of HCR labeled *ccka*+ *pitx2*+ neurons.

### Neuronal t-types are non-uniformly distributed in the tectum

To test whether t-types are spatially organized relative to the three axes of the OT volume, we selected DE genes, which were expressed in a single or a small number of clusters (Fig. 1), and examined their spatial expression patterns using multiplexed RNA in situ hybridization chain reaction (HCR). We coregistered the HCR patterns within the standard coordinates of the mapzebrain.org atlas^38,39^ and measured their expression levels at the same transverse and coronal sections (Fig. 2, Extended data Fig. 6 & 7).

We discovered that genetically identified neurons are enriched in specific domains along the SD axis. Most prominently, GABAergic and glutamatergic neurons largely populate the deepest and most superficial layer of the SPV, respectively, with cholinergic neurons sandwiched between them (Extended data Fig. 6). Individual excitatory and inhibitory t-types generally follow this rule, but can also break it. For example, the excitatory marker *bhlhe23* is expressed in deep SPV neurons, where it is surrounded by inhibitory *insm2*, *chodl, npb*, and *npy* neurons (Fig. 2, Extended data Fig. 6 & 7). The excitatory markers *ccka*, *onecut1*, *pitx2,* and *zic2*a are restricted to superficially located neurons in the SPV (Fig. 2, Extended data Fig. 6 & 7), whereas *cort*, *cckb*, and *irx1b* are expressed by neurons in the middle of the SPV (Fig. 2, Extended data Fig. 7). Some t-types are present in all SPV regions, namely those expressing *atf5b*, *sp5l,* and *neurod1*, while the inhibitory markers *esrrb* and *rpp25b* show a two-layer expression pattern (Fig. 2). We rarely found neurons expressing the same marker directly adjacent to each other (Fig. 2, Extended data Fig. 5 & 6), suggesting that neurons of the same type form a mosaic in the SPV, similar to individual RGC types in the retina.

**Fig. 6.**
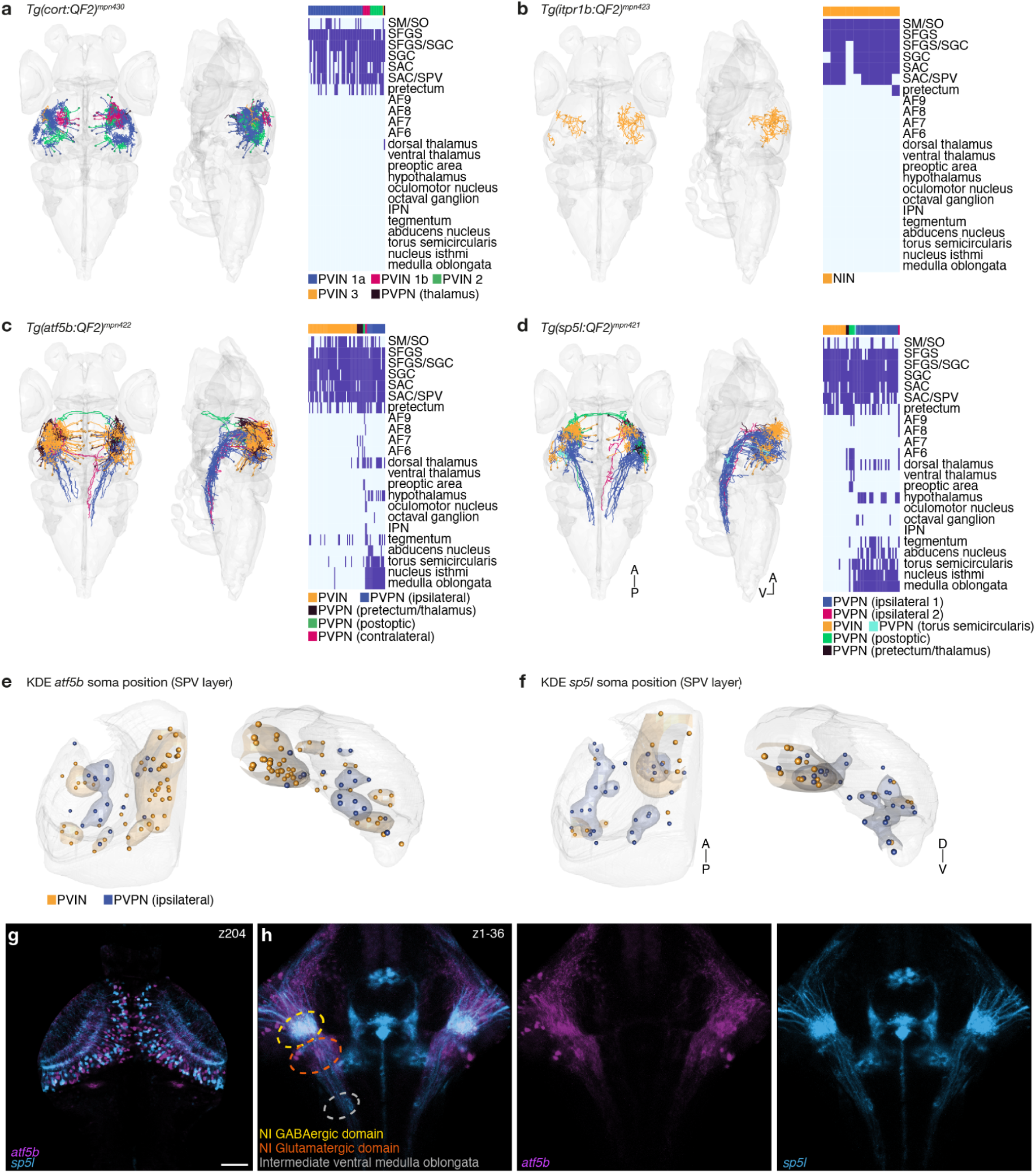
Combination of morphological types within tectal t-types. **a.** Sparsely labeled *cort+* neurons were registered into a reference brain (N=62). Right: anatomical matrix. **b.** Sparsely labeled *itpr1b+* neurons were registered into a reference brain (N=10). Right: anatomical matrix. **c.** Sparsely labeled *atf5b+* neurons were registered into a reference brain (N=77). Neurons are color coded according to their m-type. Right-anatomical matrix. **d.** Sparsely labeled *sp5l+* neurons were registered into a reference brain (N=53). Right: anatomical matrix. **e.** *atf5b* interneurons (N=49) and ipsilateral neurons (N=18) were mirrored to the left hemisphere, and a KDE was measured according to their soma position, revealing separation of m-types along the anterior-posterior and dorsal-ventral axes. Transverse and coronal views of the SPV layer are shown. **f.** *sp5l* interneurons (N=16) and ipsilateral neurons (N=29) were mirrored to the left hemisphere, and a KDE was measured according to their soma position, revealing separation of m-types along the anterior-posterior and dorsal-ventral axes. **g.** Registered confocal stack of *atf5b* and *sp5l* transgenic fish. A single focal plane spanning the OT is shown. Scale bar = 50 μm. **h.** Z-projection of the nucleus isthmi area, showing *atf5b* and *sp5l* projections forming collaterals at different parts of the nucleus isthmi. Abbreviations: A; anterior, NIN; neuropil interneurons, P; posterior, PVPN; periventricular projection neurons, PVIN; periventricular interneurons neurons, SIN; superficial interneuron, V; ventral

**Extended data Fig. 12.**
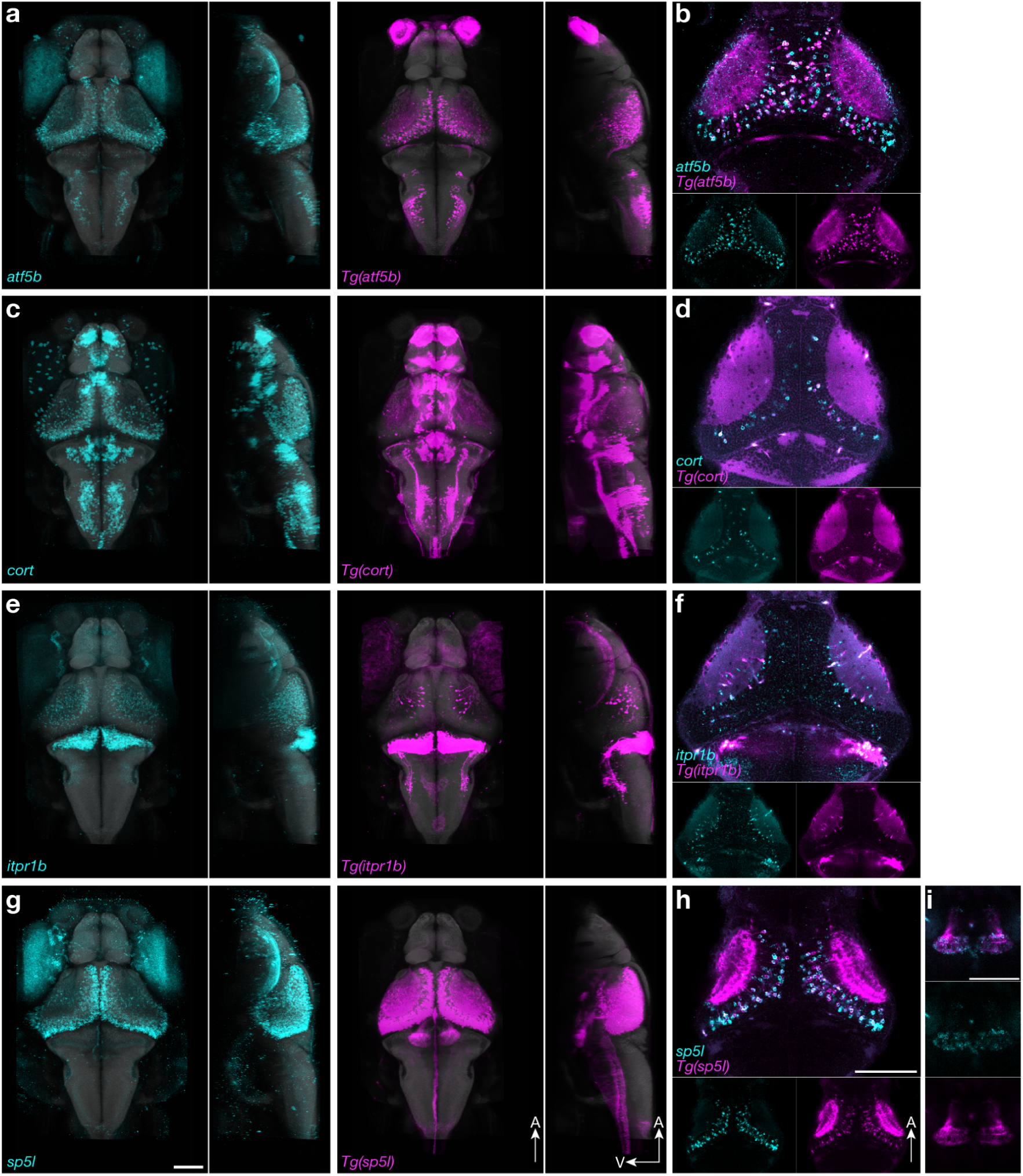
Comparison of transgenic expression with gene expression map. **a.** Average expression of *atf5b* labeled by in situ HCR (cyan) vs. knock-in transgenic (Tg) labeling of *atf5b*+ neurons (magenta). The expression pattern of 3 individual fish was registered onto the mapzebrain.org atlas and averaged. **b.** Co-labeling of *atf5b* and *Tg(atf5b)*. A single tectum plane is shown. While not all the *atf5b* detected with HCR are labeled in the transgenic line, all the transgenic labeled neurons are HCR positive. This results from either variegation of the QF2/QUAS expression, or from neurons expressing the RNA but not the protein product. **c.** Average expression of *cort* labeled by in situ HCR (cyan) vs. knock-in transgenic labeling of *cort*+ neurons (magenta). The expression pattern of 3 individual fish was registered onto the mapzebrain.org atlas and averaged. **d.** Co-labeling of *cort* and *Tg(cort)*. A single tectum plane is shown. **e.** Average expression of *itpr1b* labeled by in situ HCR (cyan) vs. knock-in transgenic labeling of *itpr1b*+ neurons (magenta). The expression pattern of 3 individual fish was registered onto the mapzebrain.org atlas and averaged. **f.** Co-labeling of *itpr1b* and *Tg(itpr1b)*. A single tectum plane is shown. **g.** Average expression of *sp5l* labeled by in situ HCR (cyan) vs. knock-in transgenic labeling of *sp5l*+ neurons (magenta). The expression pattern of 3 individual fish was registered onto the mapzebrain.org atlas and averaged. A weak expression of *sp5l* is detected in the caudal hypothalamus in comparison to the Tg line (see k). **h.** Co-labeling of *sp5l* and *Tg(sp5l)*. A single tectum plane is shown. **i.** Co-labeling of *sp5l* and *Tg(sp5l)*. A single plane of the caudal hypothalamus is shown.

**Extended data Fig. 13.**
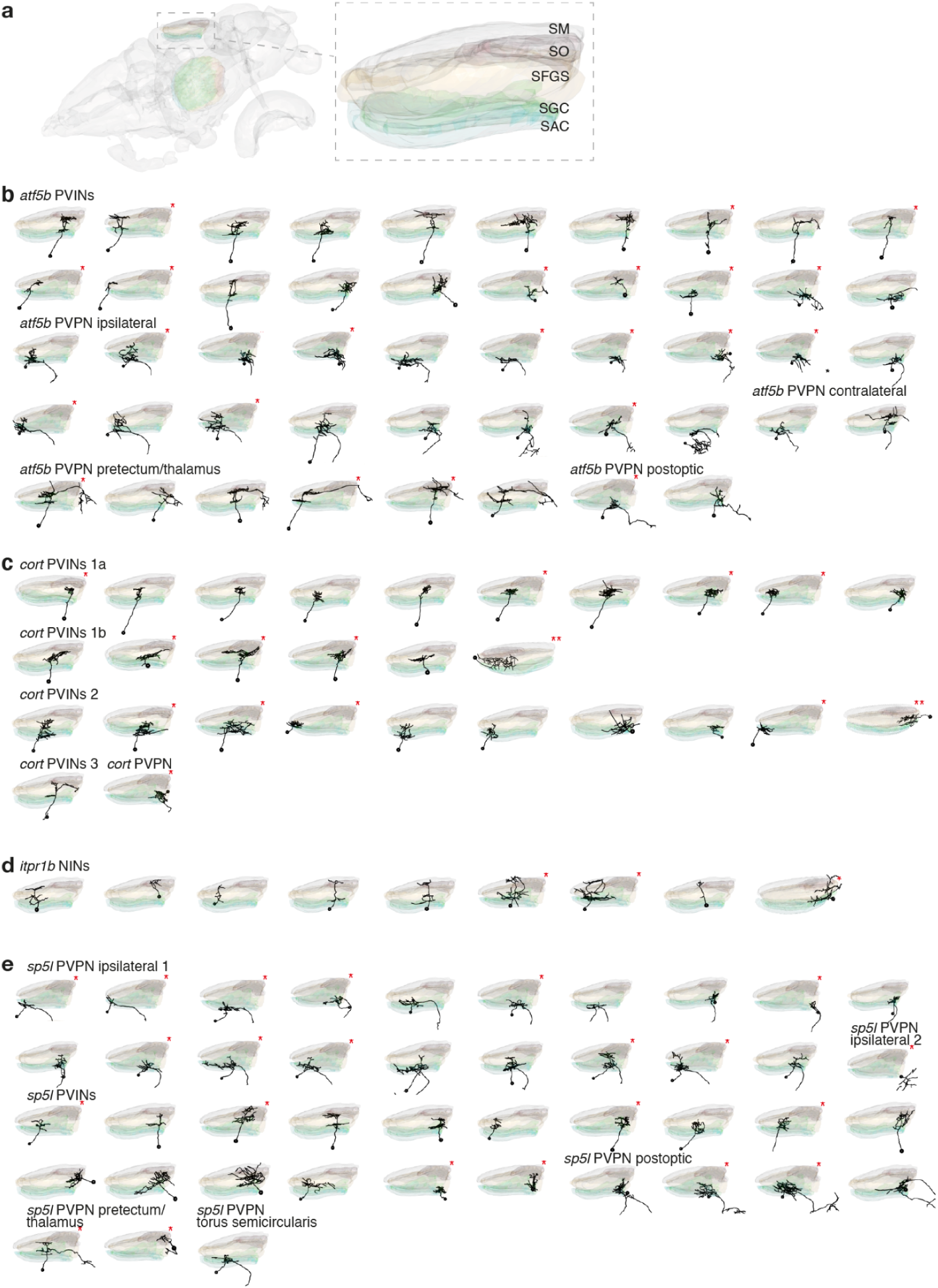
Variation of neuropil stratification patterns within t-types and m-types. **a.** Illustration of the larval zebrafish brain, highlighting the tectal neuropil layers. Abbreviations: SAC; stratum album centrale, SFGS; stratum griseum et fibrosum superficiale, SGC; stratum griseum centrale, SM; stratum marginale, SO; stratum opticum. **b. -e.** Stratification patterns of selected m-types within the tectal neuropil. * Right hemisphere located neurons, whose images were mirrored for visualization simplicity. ** lateral and not medial viewing angle.

To obtain a holistic picture of OT cell-type architecture, we manually labeled the neurons’ centroid positions (Fig. 2c-d), and measured the nearest neighbor distances (NND) in 3D, between each centroid in a given t-type to its nearest neighbors of all other t-types. We then performed hierarchical clustering on the mean NND vector for each t-type (Fig. 2e). The spatial organization of the clustered NND vectors divided the SPV into three distinct layers: superficial, intermediate, and deep (Fig. 2e-f), illustrating locally neighboring t-types. This organization disappeared with shuffled labels (Extended data Fig. 8). Given that the superficial cells of the SPV are born before the deep cells^22^, this finding suggests that, similar to the retina^40^, cell classes and cell types develop in a predetermined order, with excitatory t-types generally preceding inhibitory t-types (Fig. 2; Extended data Fig. 6c).

### Combinatorial expression patterns further separate t-types along the anatomical axes of the OT

The spatially analyzed DE genes included markers that are expressed in a single or a small number of clusters. To further examine the spatial organization of transcriptomic subtypes defined by a combination of marker genes we performed iterative multiplexed HCR labeling of submarkers for cells expressing *atf5b* and *sp5l,* respectively, and examined their spatial positioning. We focused on a combination of transcription factors and other unique effector genes (encoding mainly neuropeptides and proteins for neurotransmitter synthesis or transport^41^). *Atf5b* was strongly expressed in cluster e17 and weakly detected in several clusters, including e6, e14 and e21 (Fig. 1 & 3). Based on cluster e17 markers, we asked how *atf5b* t-types are anatomically arranged. We found that neurons co-expressing *atf5b* together with the transcription factors *foxb1a* or *etv1* were separated along the SD axis similar to the molecular layers, while co-expression with *cebpa*, which is an inhibitor of cell proliferation^42^ had higher density in the superficial posterior zone (Fig. 3a-b). Similarly, neurons co-expressing *sp5l* with the effector genes *uts1* and *chata* were separated within the intermediate molecular layer, while co-expression with the neurogenic factor *neurod1*^43,44^ was concentrated in the posterior zone (Fig. 3d-e). Overall, we find that clusters defined by either a single marker or a combinatorial code were spatially segregated along the SD axis, while distribution along the SPV anterior-posterior axis is related to neurogenic factors and the progression of neuronal differentiation from the posterior marginal proliferative zone^23,24^.

### Individual tectal t-types are selectively tuned to multiple visual stimuli

We next asked if neurons of a given t-type shared selectivity to one or multiple sensory or motor features. To this end, we recorded calcium activity by 2-photon volumetric imaging in tectal neurons of *elavl3:H2B-GCaMP6s* transgenic larvae, exposed to a sequence of behaviorally relevant visual stimuli^5,45^. The stimuli included both local and global motion cues: dots moving in different directions, either continuously or in a saltatory fashion (distinguishing a neutral, or prey-like, from a social cue), a forward moving grating (evoking optomotor responses), a looming black disk (simulating a predator or an object on a collision course), and ambient luminance changes (Fig. 4a). To determine the t-type for each of the recorded neurons, we subsequently performed iterative multiplexed HCR labeling of up to 6 mutually exclusive marker genes (except for *pitx2* and *ccka*). By co-registering the two volumetric data sets, we could unambiguously assign a t-type to 1,204 functionally characterized neurons from 6 animals (Fig. 4b, see methods for details).

We found that average t-type responses had elevated scores for at least two stimuli of the set compared to the population mean response (Fig. 4c). For example, moving dots as well as moving gratings evoked strong activity in the *atf5b* type, while *itpr1b* neurons scored highest for looming and OFF ramp stimuli (Fig. 4c). To test whether individual neurons within the same t-type or between t-types have similar functional responses we correlated calcium traces of all t-type+ neurons with each other across animals (Fig. 4d). A correlation matrix of raw calcium traces, sorted by t-type, and within t-type by overall response score to local motion stimuli, revealed stereotyped clusters of positive and negative pairwise correlations within and between all t-types (Fig. 4d). This indicates a functional diversity within each t-type, varying mostly in the relative frequency of local- and global-motion tuned neurons. Neurons co-expressing *pitx2* and *ccka*, although identified in only 12 neurons, showed a mixed functional tuning for local- and global-motion, similarly to neurons characterized by a single marker gene (Extended data Fig. 11e). Mean pairwise correlations of calcium responses were significantly higher within t-type than between t-types for six out of nine marker genes tested (Fig. 4e). We conclude that neurons of the same t-type are not coherent in their visual tuning but are, on average, functionally more similar when compared to other t-types. In line with our findings, distinct t-types in the mammalian superior colliculus, the analogous brain region to OT, also showed heterogeneous specificity in their visual responses^46^.

### Functional diversity among genetically similar neurons is spatially organized

Next, we asked whether any functional response type (f-type) in the OT was linked to one specific t-type. To this end, we first classified functional responses by clustering the highest-responding tectal neurons to each stimulus, excluding HCR labeled neurons (n=7094). This resulted in 17 distinct f-types (Fig. 5a). The response vectors formed five superclusters in a UMAP embedding, corresponding to broad classes of neurons tuned to local motion, ON, OFF, looming, and grating motion, respectively (Fig. 5b-c). We then assigned one of the 17 f-types to each HCR labeled and functionally profiled ROI (Extended data Fig. 10). The resulting distribution of f-types within t-types revealed that no f-type was specific to or strongly overrepresented in one t-type. In parallel, we investigated whether any t-type is overrepresented in a functional cluster or supercluster. (Fig. 5d, Extended data Figure 11). Transforming all HCR labeled functional ROIs into the UMAP embedding revealed that specific t-types are locally enriched within f-type superclusters. A similar result was obtained when mapping response vectors from neurons in transgenic lines expressing GCaMP6s under the control of the *atf5b*, *cort*, *itpr1b,* or *sp5l* regulatory regions (see Methods; Fig. 5e; Extended data Fig. 11), demonstrating the validity of the HCR labeling approach.

We speculated that the observed diversity of discrete f-types within t-types had its origin in the anatomical localization of individual cell bodies. To test this hypothesis, we grouped recorded neurons of the same t-type and same f-type into t/f-clusters and determined their anatomical distributions. The t/f-clusters within t-types were often separated along the AP anatomical axis: For instance, f-types 4 and 9 were almost completely non-overlapping in *itpr1b+* neurons (Fig. 5f). To quantify this observation, we computed the gaussian kernel density estimate (KDE) of anatomical cell body distributions for each t/f-cluster and measured the pairwise spatial overlap of t/f-cluster KDEs within t-types and f-types, respectively. We discovered that t/f-clusters show larger anatomical separation than shuffled controls (Fig. 5g), indicating that cell body position within a tectal t-type strongly influences its functional phenotype. Interestingly, t/f-clusters of the same f-types that are separated by t-type also show significantly lower spatial overlap within some individual f-types and across f-types.

Finally, we wondered whether the cell body location of an OT neuron is more informative of t-type identity than its functional properties. For this, we performed support vector machine (SVM) classification on t-type based on the nine marker genes, given either the anatomical cell-body position or position in the 8-dimensional functional space as input. In line with the dependency of functional identity on anatomical localization, we found that cell body position is a better statistical predictor for t-type identity than position in functional space. Combining both spaces yields even higher predictive power (Fig. 5h-i). Taken together, these results indicate a strong contribution of anatomical position to the functional phenotype of neurons with a given transcriptome.

### Transcriptome and position influence tectal neuron morphology

Tectal neurons exhibit a rich diversity of morphologies, including specific dendritic and axonal targets^5,6,47–50^. To explore whether t-types are related to the different morphological types, we sparsely labeled single neurons in transgenic reporter lines labeling the *atf5b*, *cort*, *itpr1b,* and *sp5l* t-types (Extended data Fig. 12), and traced their morphologies (see Methods). The vast majority of *cort* neurons are inhibitory periventricular interneurons (PVIN), with cell bodies located in the SPV layer. *Cort* interneurons varied in their stratification patterns: two types were monostratified within the SFGS layer, showing narrow (PVIN 1a) or wide stratification patterns (PVIN 1b), while a third type bistratified in the SGC/SAC and SFGS layers (PVIN 2). We also identified a single *cort* neuron projecting towards the dorsal thalamus, which could represent a rare morphotype (Fig 6a, Extended data Fig. 13c). The *itpr1b* neurons reside exclusively in the neuropil (Fig. 6b). Their morphologies are reminiscent of tectal tristratified pyramidal/type I neurons^47,50^, yet additional stratification patterns were also observed in this t-type, including monostratified and bistratified neurons (Extended data Fig. 13d).

The excitatory t-types *atf5b* and *sp5l* included specific sets of ipsilaterally or contralaterally projecting neurons and PVIN with a range of dendritic stratification patterns (Fig. 6c,d; Extended data Fig. 13b,e). Overall we observed that for both single marker genes (*cort*, *itpr1b*) and genes expressed in a few clusters (*atf5b*, *sp5l*), the variation in both the projection and tectal stratification exceeds the variation that can be explained by the t-type, indicating the influence of extrinsic factors on morphological differentiation.

We asked if we could further differentiate neurons with comparable projection patterns according to t-type. Registration of *atf5b* and *sp5l* confocal images highlighted the differences between their ipsilateral projections (Figure 6g-h): *sp5l* neurons have collaterals within the GABAergic domain of a tegmental nucleus, the nucleus isthmi^51^, while *atf5b* neurons form collaterals in its glutamatergic domain (Figure 6h). As is the case for t/f-clusters, the morphology of neurons belonging to the same t-type varies across the OT: *atf5b* and *sp5l* cells tend to develop into interneurons in the anterior OT and into projection neurons in the posterior OT (Fig. 6e-f). Taken together, this suggests that individual t-types accommodate a distinct range of morphologies, arranged in a position-dependent manner across the AP axis of the OT.

## Discussion

A neuron’s shape, synaptic connectivity and function, i.e., its phenotype, are dictated, or at least bounded, by the genes it expresses. High-throughput barcoding and sequencing efforts promise to offer deep insights into brain function, provided that transcriptomic datasets can be read out at scale and eventually translated into wiring diagrams^52^. This formidable task is aided by the observation that single-cell transcriptomes can be clustered by similarity of their gene expression profiles, with clusters corresponding to individual cell types. Such dimensionality reduction allows researchers to focus on characterizing the connectivity and functional properties of reproducible sets of neurons, ‘cell type by cell type’ and across animals. Single-cell RNA sequencing and spatial transcriptomics have successfully matched cell types to specific brain regions^53–55^, to developmental trajectories^56–60^ and to specific functions^2,4,7,61^, although the latter with less stringency and consistency^16,17,62^. However, there are at least two factors that confound this straightforward cell-type concept: position in the tissue and development.

First, transcriptionally similar neurons may differ locally in their phenotype in brain areas that are regionally specialized, such as topographic maps, which are prevalent in the visual, auditory, and somatosensory systems. For the retinotopically mapped OT, we find that transcriptomic cell types have overlapping sets of visual responses and morphologies, and these are organized along the spatial coordinates of the OT. We speculate that these regional differences in f-type distribution reflect their input and output heterogeneities. Such heterogeneities are well known and ethologically relevant because visual processing is adapted to stimulus statistics that are not uniform across the visual field Förster et al. 2020; Baden et al. 2020; Zhou et al. 2020).

Second, neurons are not manufactured in one sweep like transistors are printed on a microchip; they grow in number by cell division and wire up by extending neurites, often over long distances, and with their growth cones searching for molecularly matching synaptic partners. Later born neurons of the OT, which reside closer to the posterior marginal zone, are more likely to receive retinotopic information from the nasal retina. These neurons have a different set of possible synaptic partners than similar t-types born earlier and residing at the anterior OT, ultimately resulting in an asymmetric circuit layout. Additionally, synaptic partners are defined by expression of cell-surface and axon guidance molecules, and these are transiently expressed during development^60,65^, increasing the phenotypic diversity within transcriptomically indistinguishable types born later.

While our data only represent a single developmental time point, the spatial position can serve as a proxy for the neuronal time of birth. Along the superficial-to-deep axis of the tectum, the transcriptomic layering that we discovered correlates strongly with time of birth: early-born neurons come to reside closer to the surface of the SPV, as they are displaced by later-born neurons, which stay near the deep ventricular zone^21,22^. A similar temporally staggered development of cell types has been reported in vertebrate and invertebrate neuronal systems^40,66^. Thus, neurons are under the influence of dynamically changing local cues depending on when and where in the tissue they are born, differentiate and form connections.

An influence of birthdate and positional information on the expression of cell-type traits supports the notion that a limited number of global transcriptomic states are re-used and locally adapted to participate in specialized circuits and display divergent functional response profiles^67^. In this revised concept, cell types correspond to t-types. While the repertoire of genetically distinguishable t-types is already enormous^7,12,13,68^, local extrinsic cues can generate even more phenotypic variants of genetically programmed circuitry.

## Methods

### Zebrafish husbandry and maintenance

Adult zebrafish were kept at 28°C under a day/night cycle of 14/10 hours, pH of 7-7.5, and a conductivity of 600µS. The following transgenic lines were used in this study: Wild type (WT) fish of the TL stain, *Tg(elavl3:H2b-GCaMP6s)jf5, Tg(atf5b:QF2)mpn422, Tg(cort:QF2)mpn430 Tg(itpr1b:QF2)mpn423, Tg(sp5l:QF2)mpn421, Tg(QUAS:GCaMP6s)mpn164.* All larvae produced by natural matings and raised until 6 days post fertilization (dpf) at 28°C in Danieaús solution in petri dishes. The animal experiments were performed under the regulations of the Max Planck Society and the regional government of Upper Bavaria (Regierung von Oberbayern), approved protocols: ROB-55.2-2532.Vet_03-19-86, ROB-55.2-2532.Vet_02-19-16, ROB-55.2-2532.Vet_02-21-93.

### Single-cell RNA sequencing

#### Tissue dissections and cell dissociation

6-7 dpf WT larvae were anesthetized in tricaine (one larvae at a time), followed by removal of the eye and the skin around the brain using a fine Tungsten needle. The optic tectum (OT) and torus longitudinalis (TL) were carefully dissected under a stereoscope and transferred into a 1.5ml low-binding protein tube containing ∼50 µL of PBS, and placed on ice. Cell dissociation procedure was performed immediately after the dissections (see below). Eleven dissection batches were performed on different days, with 20-25 OTs and TLs dissected during each batch.

#### Cell dissociation

Cell dissociation was performed using papain (Papain dissociation system, Worthington Biochemical Corporation). A 500μl of oxygenated papain was added to the 1.5ml tube containing the 50μl PBS and the dissected tissue, and incubated at 37°C with superficial oxygen flow. The samples were gently pipetted every 10 minutes, and were fully dissociated after 45 minutes (a 2μl sample was visually inspected to assess cell separation). The cells were then centrifuged for 10 minutes at 400g, at 4°C, using a swinging bucket centrifuge. The supernatants were carefully removed, and the cells were resuspended in 260μl of equilibrated ovomucoid solution and 30 μl of DNaseI. The cells were then centrifuged for 10 minutes at 400g, at 4°C, the supernatants were removed, and the cells were resuspended in 500μl PBS and 0.04% BSA. This step was repeated once again, after which the cells were filtered using 30 μm cell strainer. Lastly, the cells were centrifuged, and the supernatants were removed, leaving 80 μl in which the cells were pipette gently. 5 μl of the dissociated cells were transferred into a new 1.5ml low-binding protein tube and mixed with 13 μl PBS, 0.04% BSA and 0.4% Trypan blue for cell counting and viability assessment.

#### Library preparation and sequencing

The dissociated cells were loaded onto a commercially available droplet-based single-cell barcoding system (10x Chromium Controller, 10x Genomics). The Chromium Single Cell 3′ Reagent Kit v3 (10x Genomics) was used to prepare single-cell 3′ barcoded cDNA and Illumina-ready sequencing libraries according to the manufacturer’s instructions. The cDNA libraries were sequenced using an Illumina HiSeq 2500 machine (batches 1-8) or Illumina NovaSeq 6000 (batches 9-11), with a mean of ∼76,000 reads per cell over the 11 batches. The raw sequencing data will be made available upon publication through NCBI’s Gene Expression Omnibus (GEO).

#### Quality check, batch correction and clustering analysis

The sequenced data was processed using CellRanger-7.1.0 (filtering, barcode and UMI counting) with default command line options. The sequenced reads were aligned to the zebrafish GRCz11 genome assembly (Ensembl release 98). To prevent confounding effects of the tissue dissection and cell dissociation on gene expression analysis, immediate early genes (IEGs) that were induced during the procedure^69^, were removed prior to the analysis. We have curated a list of 43 zebrafish IEGs (Extended data table 2), and excluded them from the cell-gene matrix. The data from each of the 11 batches was filtered using the *Seurat* R package version 5.1.0^70^, to ensure analysis of high quality cells, including filtering out cells expressing less than 600 or higher than 4000 genes, cells with higher than 7500 unique molecular identifiers (UMIs), or containing higher than 10% mitochondrial genes. Additionally, to eliminate contamination from adjacent tissues, cells expressing the habenula marker genes *gng8* and *kiss1*, as well as cells expressing the lens crystallin gene *crygm2d13* above 0.5, were excluded. The data was then normalized using the Seurat “LogNormalize” methods with a scale factor of 10,000. A set of 2,000 variable features were identified using the “vst” selection method, and the data was scaled using the “ScaleData” command. The detected variable features were used for principal component analysis. In order to filter out doublet cells from the analysis, initial clustering was performed on each batch separately, and the clustering information was used as an input for DoubletFinder^71^. Finally, the 11 batches were merged into a single Seurat object, and cells identified as doublets by DoubletFinder (n=5,931) were filtered out. The resulting dataset contained 45,766 cells with a mean of 1,224 genes per cell and 2,420 UMIs. We then performed batch correction using *Harmony*^28^, grouping the variables according to their original batch followed by dimensionality reduction with Uniform Manifold Approximation and Projection (UMAP)^72^. Clustering analysis was performed by applying the Seurat “FindNeighbors” command with reduction=”harmony” and dims=25 and the “FindCluster” command with resolution set to 0.6 (sub-clustering resolution was: neurons=1, excitatory neurons= 1.2, inhibitory neurons=1.2). The clustering resolutions were defined using clustering trees generated with the R package *Clustree*, together with SC3 stability scores^73,74^. Following the separation of excitatory and inhibitory cells, and prior to their respective clustering, cells expressing *gad1b* above 0.8 were filtered out from the excitatory dataset, while cells expressing either s*lc17a6a (vglut2b)* or *slc17a6b (vglut2a)* above 0.8 were filtered out from the inhibitory dataset. After manual inspection of marker gene expression, clusters e24, e28 and e31 were split from a single cluster, as well as clusters i3 and i16, using the “FindSubCluster” command with resolutions of 0.3 and 0.25, respectively. To evaluate clustering correlation, genes were ranked according to their specificity score, calculated using the *genesorteR* R package^75^, and correlation matrices were generated according to the top 10 scored genes per cluster. The vast majority of clusters showed low correlation with other clusters, representing unique t-types. Cluster robustness was evaluated by measuring pairwise Jaccard index of cells grouped together under different clustering parameters using the R package *scclusteval*^76^. Additionally, cluster stability was evaluated by bootstrapping as described in Tang et al. 2021^76^. 80% of the cells were sampled and reclustered. The highest Jaccard index was recorded for matching clusters for each subsample. This procedure was repeated 100 times and plotted as a Jaccard Raincloud plot. In most instances, we observed that clusters with small numbers of cells were scored low, as a result of infrequent sampling of those cells.

Generally, we took a conservative approach when defining and including clusters, which were then followed by validation of marker genes in situ (see below).

Finally, differentially expressed genes were identified using the Wilcoxon Rank Sum test integrated in the Seurat “FindAllMarkers” command. Pseudotime analysis was performed using Monocle3^77^.

The full R analysis code will be made publicly available upon publication.

### Multiplexed and iterative *in situ* hybridization chain reaction (HCR)

The HCR experiments were performed on *Tg(elavl3:H2b-GCaMP6s)jf5* (to enable registration onto a common coordinate system) or the transgenic knock-in larvae labeling *atf5b*, *cort*, *itpr1b*, and *sp5l* expressing cells. All the HCR reagents including probes, hairpins and buffers were purchased from Molecular Instruments (Los Angeles, California, USA). The staining was performed according to previously published and modified protocol^39,78^. The larvae were anesthetized in 1.5 mM tricaine and fixed with ice-cold 4% PFA/DPBS overnight at 4°C with gentle shaking. The following day, larvae were washed 3 times for 5 minutes with DPBST (1x Dulbecco’s PBS + 0.1% Tween-20) to stop fixation, followed by a short 10-minute treatment with ice-cold 100% Methanol at −20°C to dehydrate and permeabilize the tissue samples. Next, rehydration was performed by serial washing of 50% MeOH/50% DPBST and 25% MeOH/75% DPBST for 5 minutes each and finally 5 × 5 minutes in DPBST. 10-12 larvae were transferred into a 1.5 ml tube and pre-hybridized with pre-warmed hybridization buffer for 30 minutes at 37°C. Probe solution was prepared by transferring 2 pmol of each HCR probe set (2 µl of 1 µM stock) to 500 µl of hybridization buffer at 37°C. The hybridization buffer was replaced with probe solution, and the samples were incubated for 12-16 hours at 37°C with gentle shaking. To remove excess probes, larvae were washed 4 × 15 minutes with 500 µl of pre-warmed probe wash buffer at 37°C. Subsequently, larvae were washed 2 × 5 minutes with 5x SSCT (5x sodium chloride sodium citrate + 0.1% Tween-20) buffer at room temperature. Next, pre-amplification was performed by incubating the samples in 500 µl of amplification buffer for 30 minutes at room temperature. Separately, 30 pmol of hairpin h1 and 30 pmol of hairpin h2 were prepared by snap-cooling 10 µl of 3 µM stock by incubating the hairpins in 95°C for 90 seconds, and cooling down to room temperature in a dark environment. After cooling down for 30 minutes, hairpin solution was prepared by transferring the h1 and h2 hairpins to a 500 µl amplification buffer. The pre-amplification buffer was removed and the samples were incubated in the hairpin solution for 12-16 hours in the dark at room temperature. Excess hairpins were washed the next day 3 × 20 minutes using 5x SSCT at room temperature. Larvae were then long-term stored at 4°C in 5X SSCT until imaging.

For iterative HCR staining, the HCR probes and hairpins were stripped using DNAseI treatment. The imaged fish were placed separately in 1.5ml tubes and incubated in a mix of 5µl 10x reaction buffer (Invitrogen #AM2238), 5µl Turbo DNAseI (final concentration 0.2U/µl, Invitrogen #AM2238) and 40µl DPBS for 4 hours at 37°C. The samples were then washed 3 × 5 minutes with DPBST and the complete removal of the HCR signal was validated under a confocal microscope. The HCR striped fish were kept separated in 1.5ml tubes and underwent another round of HCR staining and imaging, beginning from the pre-hybridization step.

HCR data was registered onto the HCR reference brain as previously described^39^ using Advanced Normalization Tools (ANTs)^79^. For the functional HCR experiments (see below), where iterative HCR stainings were performed, each iteration was registered to the first iteration of the same animal. A total of 3 HCR rounds were performed on single animals without any noticeable morphological distortions or reduction of the endogenous fluorescence of the *Tg(elavl3:H2b-GCaMP6s)* used as the reference registration channel.

### HCR image registration

All the HCR and immunostaining images were aligned onto the mapzebrain.org *Tg(elavl3:H2b-GCaMP6s)* average brain^38,39^.

The ANTs registration command used was: antsRegistration -d 3 --float 1 -o [${output1},${output2}] -- interpolation WelchWindowedSinc --use-histogram-matching 0 -r [${template},${input1},1] -t rigid[0.1] -m MI[${template},${input1},1,32,Regular,0.25] -c [200×200×200×0,1e-8,10] -- shrink-factors 12×8×4×2 --smoothing-sigmas 4×3×2×1vox -t Affine[0.1] -m MI[${template},${input1},1,32,Regular,0.25] -c [200×200×200×0,1e-8,10] -- shrink-factors 12×8×4×2 --smoothing-sigmas 4×3×2×1 -t SyN[0.01,6,0.0] -m CC[${template},${input1},1,2] -c [200×200×200×200×10,1e-7,10] --shrink-factors 12×8×4×2×1 --smoothing-sigmas 4×3×2×1×0

followed by applying the transformation files on the HCR image channels using the ANTs command: antsApplyTransforms -d 3 -v 0 -- float -n WelchWindowedSinc -i ${input3} -r ${template} -o ${output4} -t ${output1}1Warp.nii.gz -t ${output1}0GenericAffine.mat

All the registered HCR data used in this study are publicly available through the mapzebrain.org atlas.

### Confocal imaging

HCR labeled samples were embedded in 2% low-melting agarose in 1x DPBS (Dulbecco’s PBS) and imaged with a Zeiss LSM700 confocal scanning microscope (upright), equipped with a 20x water immersion objective. Z-stacks, composing 2 tiles (or 1 tile for functional HCR experiment), were taken and stitched to produce a final image with size of 1039 × 1931 pixel (463.97 × 862.29 µm, 1 µm in z).

### Pixel intensity quantification of HCR data

For each HCR labeled gene, three larvae were imaged and registered as described above. A maximum intensity projection was generated for each image for planes z289-z291. Coronal sections were generated using Fiji “reslice” option^80^, and the maximum intensity projection was generated for planes x463-465. ROIs spanning the SPV for both the transverse and coronal section were used to measure the pixel intensity profile using a custom Fiji macro. The data was analyzed and plotted using R.

### HCR positive cells labeling

Cell centroids in the HCR data were manually labeled using napari points layer tool^81^, by examination of HCR signal together with the nuclear labeling of the *Tg(elavl3:H2b-GCaMP6s)*. For identifying co-labeled submarkers in *atf5b* and *sp5l* expressing cells, polygons were first manually drawn using napari around the cells expressing the main markers. The expression of submarkers were manually determined by labeling positive centroids within the main polygons. The centroids were then registered onto a reference brain using ANTs and the R package ANTsR. To define the molecular layers of the OT, the nearest-neighbor distances between centroids were measured in 3D using the R package spatstat^82^. The centroids were visualized using the R package Plotly, hierarchically clustered using complete linkage method implemented in the “hclust” function with default parameters, and plotted using the R package pheatmap^83^.

### Functional two-photon calcium imaging of *Tg(elavl3:H2b-GCaMP6s)*

Two-photon functional calcium imaging was performed on 6–8 dpf *Tg(elavl3:H2b-GCaMP6s)* larvae that expresses a nuclear calcium indicator in all neurons, without paralysis or anesthesia. The animals were embedded in 2% agarose and mounted onto the stage of a modified two-photon moveable objective microscope (MOM, Sutter Instrument, with resonant-galvo scanhead) with a ×20 objective (Olympus XLUMPLFLN, NA 1.0) and recorded for 20 min. Fish that drifted along the dorsoventral axis in the preparation were excluded from analysis. Volumetric imaging of the tectum was performed with a custom-built remote focusing arm ^45^. Refocusing through the remote arm enabled rapid sequential imaging of 6 planes (512×512 pixels) spanning 60-100 µm of the tectum along the dorsoventral axis at 5 volumes per second. In each fish neuronal activity in the tectum was recorded over two 10-minute sessions to cover the whole tectum, resulting in 12 imaging planes per fish in total. Laser power out of the objective ranged from 10 mW to 15 mW.

### Two-photon functional calcium imaging of transgenic lines

Animals were embedded and placed under a commercial two-photon laser scanning microscope (Femtonics) or custom-built two-photon microscope as previously described^45,84^. Sequential single-plane imaging at either 5 fps (custom-built microscope) or between 1-1.5 fps (Femtonics) was performed at 4-10 different depths along the dorsoventral axis of the tectum for a 10-minute session each, with the same stimulus set as detailed below. No anatomical stack (see below) was acquired from these animals, hence no registration was performed.

### Visual stimulation

Visual stimuli were designed with PsychoPy and projected by an LED projector (Texas Instruments, DLP Lightcrafter 4500, with 561 nm long-pass filter) from below onto Rosco tough rolux 3000 diffusive film water-immersed in a 10 cm petri dish. The embedded animal was placed into a 6 cm petri dish on top of the diffusive paper in the larger dish, with a spacer in between that ensured a water film between the diffusive paper and the smaller petri dish. The fish was placed at 12 mm distance from the projection screen. The fish head was manually centered in the imaging chamber, aided by projecting crosshairs on the screen. Stimuli were shown in a predetermined sequence: gratings, dot motion (8x, see below for details), gratings, OFF ramp, ON ramp, looming (2x). All stimuli were shown black on a red background to not interfere with the green GCaMP6s fluorescence signal. Individual stimulus presentations were separated by 20 seconds inter-trial intervals, except for the two loom stimuli, for which a 1-min interval was used. The whole duration of the stimulus protocol was 10 minutes.

#### Moving gratings

Gratings moving caudo-rostrally with respect to the fish were shown once at the beginning and after the dot stimuli with 20 mm width and 2 Hz temporal frequency for 20 seconds.

#### Dot motion

A black dot moving on a circular trajectory (radius, 18 mm) was shown starting in front of the fish. The dot stimulus moved either in discrete jumps at 1.5 Hz or perceptually smooth at 60.0 Hz (projector frame rate) with the same overall speed of 5 mm s^−1^ (15.9 degrees (deg) s^−1^), resulting in a stimulus duration of 22.6 seconds. Each frequency was presented using a dot diameter of 4 mm (12.7 deg). Both clockwise and counter-clockwise presentations were shown, resulting in 4 different stimuli. The whole dot stimulus set was repeated twice in each session, so 8 dot stimuli were shown in succession.

#### OFF/ON ramp

Whole-field luminance of the projected blank image was decreased to zero over the course of 2 seconds, and ramped up to normal background luminance within 2 seconds after 20 seconds delay.

#### Looming

An expanding disk was displayed with expansion from 0.6 deg to 110 deg in 83 ms centered below the fish.

### z-Stack acquisition, registration and ROI matching

After each functional recording, a high-resolution 2-photon *z*-stack of large parts of the brain including the full midbrain region was taken (1,024 × 1,024 pixels, 1 µm in *z*, 835 nm laser wavelength, plane averaging 100×). Each time series average of the 12 imaging planes were registered to this stack using the *scikit-learn* template matching algorithm. The 2-photon brain volume was then registered to the volume of the first round HCR confocal imaging as described above (see **HCR image registration**). To transform functional ROIs from 2-photon space into HCR space, ROI pixel coordinates were transformed first from imaging plane reference frame (RF) to 2-photon z-stack RF and finally to the first round HCR RF the by running the ANTs command *antsApplyTransformsToPoints* two times using the respective transformation matrices from each registration step. HCR cell centroid annotations from each round were transformed into the first round HCR RF, and all coordinates were used as seeds for generating 3×3 pixel volumes that were overlaid with registered functional ROI pixels. HCR centroids were assigned to a functional ROI based on the largest fractional overlap. Finally, all assigned as well as unassigned functional ROIs were transformed to the average brain as described above.

### Data analysis for two-photon imaging

*Suite2p*^85^ was used for motion correction, ROI detection, ROI classification, and signal extraction (time constant *tau*=7 s, diameter=4 pixels). In detail, raw volume recording files were deinterleaved into individual time series for each imaging plane. Rigid and non-rigid motion correction was performed with *suite2p* on a low-pass filtered time series in xy (gaussian, sigma=4). The motion-correction shifts were applied to the raw imaging time series. ROIs were detected on fivefold downsampled & motion-corrected time series and fluorescent traces were extracted using the average pixel intensities of ROIs over time. All functional ROIs were initially thresholded based on built-in *suite2p* classification algorithm *iscell* and an anatomical tectal mask drawn in the reference brain (see mapzebrain.org).

#### Response score

For each stimulus, a regressor was constructed by convolving a boxcar function with an exponential decay kernel that mimics the H2B::GCaMP6s off-kinetics. The resulting 8 regressors were separately fitted to the calcium trace of each functional ROI, using a linear regression model on the stimulus time window. The response score was calculated as the product of the regression coefficient (equivalent to dF/F) and the coefficient of determination R^2^. For clustering and dimensionality reduction (Fig 5), the analysis included all functional ROIs that scored above the 95th percentile of the population response score for any of the visual stimuli. Response score vectors of all high-scoring neurons excluding any t-type positive ROIs (n=7094) were scaled to unit variance and zero mean of the overall tectal population for subsequent analysis.

#### Hierarchical clustering & embedding

Functional clusters were identified in a two-step analysis: First, exemplars of the 7094 high-scoring ROIs in 8-dimensional functional space were identified using affinity propagation (*sklearn.cluster.AffinityPropagation*, default parameters). Exemplars with less than 10 associated ROIs were excluded from subsequent analysis. The remaining 169 exemplars were clustered using hierarchical clustering (*scipy.cluster.hierarchy.linkage*, method=’complete’, metric=’correlation’; *scipy.cluster.hierarchy.fcluster,* criterion=’maxclust’). In parallel, the 2-dimensional embedding for visualizing functional space was computed from the 8-dimensional response vectors of all high-scoring ROIs using UMAP (*n_neighbors=25, min_dist=0.7, metric=’euclidean’,*). Functional ROIs matched with HCR in situ labels or functional ROIs from transgenic lines were transformed into the initial UMAP space by applying the UMAP transform function to their respective response vectors.

#### Anatomical overlap metric

3D coordinates of each t/f-subcluster that contained more than 10 ROIs were used to generate a gaussian kernel density estimate with *scipy.stats.gaussian_kde(bandwidth=0.75).* For computing pairwise overlap, the KDEs were sampled, normalized, and the minimum of the joint KDEs was taken at each point. The overlap metric is bounded from 0 to 1 (0 if no overlap, near 1 if the same). Controls were generated by shuffling t-type labels of all ROIs in the whole dataset.

#### T-type classification

Response scores of t-type+ ROIs were scaled to unit variance and zero mean. A support vector machine (SVM) classifier (*sklearn.svm.SVC*, kernel=’rbf’, gamma=’scale’, C=1) was trained on 90 % of all t-type+ response vectors and gene labels. To counter the uneven distribution of t-types, the training data was upsampled to 1000 samples per t-type. Evaluation of the classifier performance was done on 10 % holdout test data. This process was repeated 20 times with permutated training/test data splits. The same classification was performed with anatomical centroid positions of t-type+ ROIs as dependent variables as well as using both response vectors and anatomical positions. Each classification was run again with shuffled marker gene labels as negative control.

### Generation of knock-in transgenic lines

The knock-in (KI) lines *Tg(sp5l:QF2)mpn421*, *Tg(atf5b:QF2)mpn423*, *Tg(itpr1b:QF2)mpn424, and Tg(cort:QF2)mpn430* were generated by locus-specific insertions using CRISPR-Cas9 and the GeneWeld approach^86^. gRNA target sequences were identified using the CCTop tool^87^. The gRNA target sequences are: *atf5b* 5’-ATTTGGACGTCATGCTCCAGAGG-3’; *cort*; 5’-GCCCCTGGAGTCCCGTCTGG-3’; *itpr1b* 5’-CATCTGCTCCCTGTATGCGGAGG-3’; *sp5l* 5’-AGGCTCGCAGCTCCCTTACGAGG-3’. Short homology sequences of 48bp spanning the upstream and downstream of gRNA site were ordered as complementary oligonucleotides (MWG) and cloned into donor plasmids using the GoldenGATEway strategy^88^. Universal gRNAs (ugRNA)^86^ were introduced into the donor to release the insert from plasmid after injection. The order of components of all the donor constructs was the following: ugRNA, upstream homology arm, short GSG linker, T2A, QF2, polyA signal, downstream homology arm and second ugRNA (inverted). CRISPR-Cas9 RNP complex was prepared at a concentration of 1.5 μM as described before^4^. The gRNA was produced by annealing customized crRNA (IDT, Alt-R® CRISPR-Cas9 crRNA) with tracrRNA (IDT, CAT# 1072533) in annealing buffer (IDT, CAT# 11-05-01-12). The gRNA was incubated with Cas9 protein (IDT, CAT# 1081060) for 15 minutes at 37°C and the donor plasmid was added to the injection mix, at a final concentration of 20 ng/μl. The CRISPR-Cas9 mix was injected into *Tg(QUAS:epNTR-RFP)mpn165* embryos at the single-cell stage. Positive transient expressor fish were raised and screened at adulthood for germline transmission.

### Cellular tracing and morphology analysis

Single neurons were sparsely labeled either during the KI generation procedure in mosaic F0 animals, or by transiently microinjecting 12.5 ng/μL QUAS:eGFP-caax plasmid into the *Tg(atf5b:QF2)mpn423, Tg(cort:QF2)mpn430, Tg(itpr1b:QF2)mpn424, or Tg(sp5l:QF2)mpn421* transgenic embryos at the single-cell stage. At 6 dpf the injected larvae were anesthetized in a lethal dose of tricaine, fixed in 4% PFA and immunostained with Mouse anti-ERK1/2 (Cell Signaling Technology) and Chicken anti-GFP (Invitrogen), according to the protocol previously published^38^.

Confocal imaging was performed as described above. Individual neurons were semi-automatically traced using the freeware NeuTube^89^, saved as SWC files and registered to the mapzebrain.org atlas using ANTs as described above and previously^38^.

The traced neurons were plotted using the R package *natverse*^90^ and manually clustered according to the projection terminals or neuropil stratification patterns. A matrix for the anatomical crossing areas for each neuron was generated with the mapzebrain.org atlas, and the heatmaps were plotted using the R package *pheatmap*^83^. Soma positions were used to generate a gaussian kernel density estimate using the R packages *misc3d* and oce^91^ and plotted using the R package *rgl*^92^.

## Code availability

All the Python, R, and ImageJ custom scripts used in this study will be made publicly available upon publication.

## Acknowledgments

We thank Marja Driessen, Rin Ho Kim and the Max Plank of Biochemistry NGS facility for their technical support. We thank Mariam Al Kassar and Nouwar Mokayes for their assistance in mapping neuronal traces to the mapzebrain.org. We thank Hagar Lavian, Ruben Portugues, and Takashi Kawashima for fruitful discussions. We thank all of the Baier lab members for their support. Funding was provided by the Max Planck Society. I.S. was supported by an Alexander von Humboldt foundation research fellowship and by the Weizmann Institute of Science Advancing Women in Science bridge position program. J.M.K. and M.S were supported by a Boehringer Ingelheim Fonds graduate fellowship. J.L. was supported by a NARSAD Young Investigator Award.

## Author contributions

I.S. performed scRNA-seq and analyzed the data. J.M.K. and I.S. performed functional two-photon imaging experiments. J.M.K. and I.S. performed registration of two-photon and HCR data. J.M.K. analyzed functional imaging data. I.S. and E.K. performed HCR staining and imaging. I.S. analyzed HCR data. J.C.D. advised on imaging data analysis. J.L. and M.W.S. performed pilot behavioral experiments. M.S. designed CRISPR constructs. I.A.A. constructed KI plasmids. I.S., E.K., E.L. and J.L. generated transgenic lines. I.S. and E.L. performed sparse labeling of neurons. E.L. immunostained, imaged, traced and registered single neuronal morphologies. I.S., J.M.K. and H.B. interpreted the data. I.S., J.M.K. and H.B. wrote the paper with input from J.L. All of the authors reviewed and edited the manuscript. H.B. supervised the project.

